# Identification of a targetable ST2-expressing fibroblast subset driving Peutz-Jeghers syndrome polyposis

**DOI:** 10.1101/2023.11.29.568817

**Authors:** Eva Domènech-Moreno, Wei-Wen Lim, Anders Brandt, Toni T Lemmetyinen, Emma W Viitala, Tomi P Mäkelä, Stuart A Cook, Saara Ollila

**Affiliations:** HiLIFE-Helsinki Institute of Life Science, University of Helsinki, Helsinki, Finland; iCAN Digital Precision Cancer Medicine Flagship, University of Helsinki, Helsinki, Finland; National Heart Research Institute Singapore, National Heart Centre Singapore, 169609, Singapore; Cardiovascular and Metabolic Disorders Program, Duke-National University of Singapore Medical School; MRC-London Institute of Medical Sciences, Hammersmith Hospital Campus, London, UK; Translational Cancer Medicine Program, University of Helsinki, Helsinki, Finland

## Abstract

Peutz-Jeghers syndrome (PJS) is associated with early-onset and recurring gastrointestinal hamartomatous polyposis caused by hereditary inactivating mutations in the tumor suppressor gene LKB1 (STK11). Due to lack of efficient prophylactic therapies PJS patients require regular surgical interventions and have an increased risk of cancer. LKB1-deficient fibroblasts have been identified as drivers of polyposis, but a safely druggable target remains to be identified. Here we investigate tumorigenic mechanisms in PJS polyps using single-cell RNA sequencing of predisposed and tumor tissue and identified a ST2-expressing crypt top fibroblast (CTF) cluster enriched in polyps. The relevance of CTFs was supported by the fast polyposis following deletion of Lkb1 in CTFs. The transcriptional signature characterizing the ST2+ fibroblasts (ST2-CTFs) was enriched in inflammation-associated fibroblasts, and PJS polyposis was exacerbated by inflammation. Cell-cell communication analysis identified the ST2-CTF signature gene interleukin 11 (IL-11) as an upstream regulator of the ST2-CTFs, and consistently, reprogramming toward ST2 fibroblasts in vitro was dependent on IL-11. Importantly, a neutralizing IL-11 antibody efficiently reduced the tumor burden in a PJS mouse model. In summary, our results reveal ST2+ tumorigenic fibroblasts as drivers of PJS polyposis and identify IL-11 as the key mediator and a potential therapeutic target in PJS.

## Introduction

Peutz-Jeghers syndrome (PJS) is a hereditary condition with an estimated prevalence of 1 in 50,000–200,000 live births (Giardiello & Trimbath, 2006). PJS is characterized by the emergence of multiple benign hamartomatous polyps throughout the GI tract and an increased risk of cancer (Jelsig et al., 2023). Regular removal of polyps using endoscopy or surgery is often required to avoid life-threatening intussusception, resulting in some patients experiencing significantly reduced quality of life. PJS is inherited through heterozygous inactivating mutations in the tumor suppressor gene *LKB1* (*STK11*) (Hemminki et al., 1998), and *Lkb1^+/-^* mice recapitulate the polyp formation with development of GI hamartomatous polyps with full penetrance, allowing mechanistic studies (Rossi et al., 2002). Previously, we and others have demonstrated that LKB1 deletion limited to the mesenchymal compartment, or specifically to fibroblasts, was sufficient for polyp formation, suggesting a key role for stromal fibroblasts in the development of PJS tumors (Cotton et al., 2022; Katajisto et al., 2008; Ollila et al., 2018). Also T-cells have been suggested as potential drivers of PJS polyposis (Poffenberger et al., 2018).

Recent work using single-cell RNA sequencing (scRNAseq) of gastrointestinal tissues has revealed the existence of spatially and functionally distinct fibroblast populations in the intestine (McCarthy, Manieri, et al., 2020; Pærregaard et al., 2023), colon (Brügger et al., 2020; Pærregaard et al., 2023) and stomach (Kim et al., 2020; Manieri et al., 2023). These include crypt top fibroblast (CTF) subsets expressing Bone morphogenic protein (BMP) ligand genes such as *Bmp2*, *Bmp5,* and *Bmp7* to support differentiation, and crypt bottom fibroblasts (CBFs) providing stemness-supporting factors such as Gremlin-1 (*Grem1*) and R-spondin 3 (*Rspo3*). GI fibroblasts promote tissue regeneration during inflammation, and abnormally and persistently activated fibroblasts are noted in fibrosis, chronic inflammatory diseases, and tumorigenesis (Wei et al., 2021). The fibroblast subsets driving PJS polyposis have not been characterized.

There have been several attempts to reduce tumor burden in PJS models. Despite some effects using long-term high-dose treatment of PJS mice with rapamycin (Robinson et al., 2009), the polyposis appears to develop independently of the AMPK-mTORC1 axis (Cotton et al., 2022; Ollila et al., 2018). COX2 expression is elevated in some polyps, and a COX2 inhibitor, celecoxib, was noted to reduce polyp burden (Udd et al., 2004), indicating importance for inflammatory pathways. We (Ollila et al., 2018) and others (Poffenberger et al., 2018) have shown that the polyps display active JAK/STAT signaling and that JAK inhibition dramatically reduces the PJS polyp burden. However, inhibition of JAK signaling is associated with significant side effects hindering long-term use. The precise mechanisms through which LKB1-deficient stromal cells promote PJS polyposis are unclear, complicating the design of targeted therapies with acceptable safety profiles.

In this study, we compare fibroblast populations in normal tissue and polyps using scRNA-seq of the normal gastric antrum and antral polyps and investigate tumorigenic mechanisms. We report a tumor-enriched fibroblast subpopulation characterized by the expression of ST2 (encoded by *Il1rl1*) and Interleukin-11 (*Il11*) driving PJS polyposis. Importantly, we show that targeting IL-11 impedes the LKB1 loss-induced fibroblast activation and efficiently reduces tumorigenesis in PJS mouse models. Our discoveries shed light on the role of stromal cells in driving the development of PJS polyps and suggest IL-11 as a potent therapeutic target for PJS-associated tumors. The findings may have implications also in a wider context to understand how fibroblast reprogramming in the tumor microenvironment can affect GI tumorigenesis.

## Results

### Enrichment of *ST2^+^* fibroblasts in PJS polyps

To get insight into the distribution and signaling activity of different cell types in PJS tumorigenesis, we performed scRNAseq of polyps and normal tissue isolated from *Twist2-Cre;Lkb1^fl/+^;R26R-tdTomato* mice. In these mice, *Lkb1* heterozygosity is restricted to mesenchymal lineages, yet they develop PJS-type polyps with similar kinetics compared to *Lkb1^+/-^*mice (Ollila et al., 2018). The polyps and adjacent normal tissue were dissected from the distal part of the stomach, the antrum, structurally similar to the colon and intestine with epithelial crypts (glands) opening to the gastric lumen, and the bottom of the crypt hosting the epithelial stem cells marked by Lgr5 (Leushacke et al., 2013). In total, we sequenced 17933 cells that, after quality control, resulted in 14641 cells representing 6112 cells from normal antrum and 8529 cells from polyps, clustering into epithelial cells (*Epcam*^+^), endothelial cells (*Pecam1*^+^), immune cells (*Ptprc*^+^), smooth muscle cells (*Hhip^+^, Myh11*^+^), pericytes (*Rgs5*^+^), and fibroblasts (*Pdgfra*^+^, *Pdpn*^+^) (Figure 1A). All cell types were found in both normal tissue and polyps (Supplementary Figure 1A-B). The *tdTomato* reporter expression, indicating recombination, was noted in mesenchymal populations but was absent in epithelial and immune cells, indicating that LKB1 loss in immune cells such as T-cells is not required for polyp formation (Supplementary Figure 1C).

**Figure 1.**
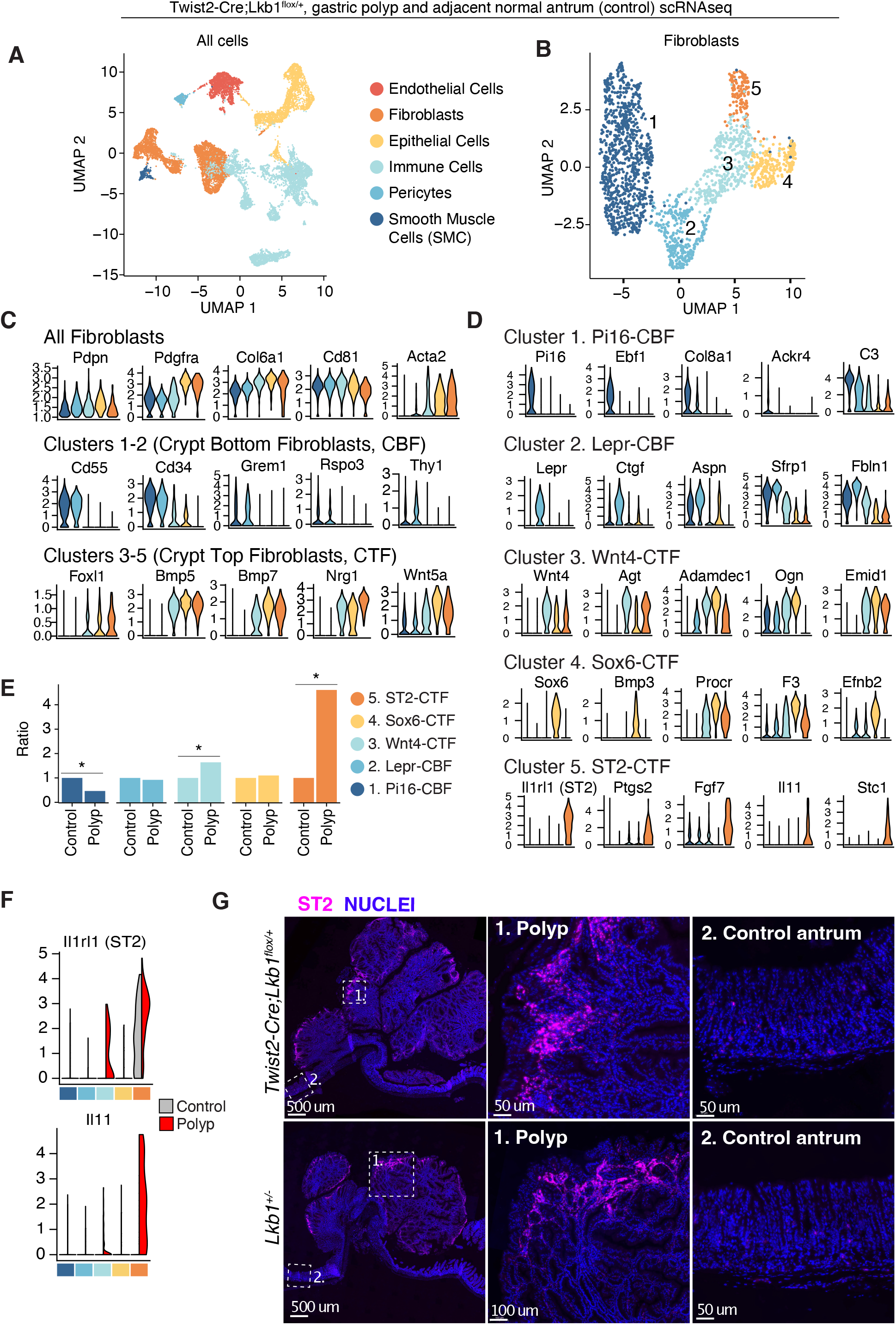
ST2-expressing fibroblasts are enriched in PJS polyps. **(A)** UMAP clustering of all cells identified in the whole scRNAseq dataset including normal stomach and gastric polyp tissue. **(B)** UMAP clustering of the fibroblast subpopulations from A. **(C)** Violin plots of markers shared by all fibroblasts as well as established markers for crypt bottom and crypt top fibroblast subsets. **(D)** Violin plots of top five markers of the identified fibroblast subpopulations. **(E)** Ratio of fibroblast subtype representation in tumors as compared to control gastric mucosa. *, p<0.05, Chi-square test. **(F)** Split violin plots indicating expression of *Il1rl1* and *Il11* expression across all fibroblast populations split by sample (grey: control, red: tumor). **(G)** Immunofluorescent staining of ST2 in tumors (T) and adjacent control mucosa (C) of *Lkb1^+/-^* or *Twist2Cre;Lkb1^fl/+^* mice.

As fibroblasts appear to represent important tumor-initiating cells in PJS polyposis (Ollila et al., 2018), the analysis initially focused on fibroblast populations. We identified five distinct fibroblast clusters positive for the pan-fibroblast markers *Pdpn*, *Pdgfra,* and *Col6a1* (Figure 1B-C). We compared these with fibroblast populations characterized in the GI tract (Brügger et al., 2020; Brügger & Basler, 2023; Manieri et al., 2023; Pærregaard et al., 2023), consisting of: i) the *Pdgfra*-low, *Cd34*^+^, *Cd55*^+^ crypt bottom fibroblasts (CBFs) forming the niche for the stem cells residing at the crypt bottom by secreting stemness-supporting factors such as R-Spondin 3 (encoded by *Rspo3*) as well as the Bmp inhibitor Gremlin 1 (encoded by *Grem1*); and ii) the *Pdgfra*-high, *Cd34*^-^, *Cd55*^-^ crypt top fibroblasts (CTFs), promoting epithelial differentiation in the upper parts of the crypts by expressing e.g. BMP ligands. In our gastric fibroblasts, clusters 1 and 2 were positive for *Cd55* and *Cd34* and expressed *Grem1* and *Rspo3* along with low levels of *Pdgfra,* suggesting that they represent CBFs. However, as opposed to colonic and intestinal fibroblasts where *Cd81* is specifically expressed in one subset of the *Pdgfra* low fibroblasts (Brügger et al., 2020; McCarthy, Manieri, et al., 2020), expression of *Cd81* was similar across all gastric fibroblast populations (Figure 1C). Instead, cluster 1 cells specifically expressed *Pi16*, *Ebf1, Col8a1*, *Ackr4,* and *C3* (Figure 1D), aligning with universal fibroblasts that mainly reside in GI submucosa (Buechler et al., 2021; Muhl et al., 2020; Thomson et al., 2018). We named these cells Pi16-CBFs. The second CBF cluster specifically expressed *Lepr* and *Ctgf* as well as high levels of *Aspn, Sfrp1*, and *Fbln1*, and we labeled them as Lepr-CBFs (Figure 1D). The characteristic CBF gene profile (Figure 1C), along with an absence of Pi16-CBFs markers, suggested Lepr-CBFs are located in the lamina propria close to the crypt/gland base as previously shown for intestinal fibroblasts expressing *Lepr* (Deng et al., 2022; Manieri et al., 2023).

CTFs in the colon and intestine express differentiation-promoting ligands such as *Bmp5*, *Bmp7, Wnt5a, Nrg1,* as well as the subepithelial telocyte marker *Foxl1 (Aoki et al., 2016; Brügger et al., 2020; Lemmetyinen et al., 2023; McCarthy, Manieri, et al., 2020; Shoshkes-Carmel et al., 2018)*. These ligands were highly expressed in our fibroblast clusters 3-5, indicating a CTF identity (Figure 1C). The cells in cluster 3 were, however, low in *Pdgfra* expression but expressed high levels of the non-canonical Wnt ligand *Wnt4* (Figure 1C-D), and we named them Wnt4-CTFs. The Wnt4-CTFs resembled a fibroblast subset that was recently shown to be subject to BMP-mediated regional regulation of niche factor expression exerted by subepithelial fibroblasts in the colon and intestine, restricting *Wnt4* expression in the upper parts of the crypts (Kraiczy et al., 2023). Cells from cluster 4 specifically expressed *Sox6* and *F3* along with higher expression levels of *Acta2,* previously identified markers for subepithelial myofibroblasts (SEMFs) corresponding to CTFs (Kinchen et al., 2018; McCarthy, Kraiczy, et al., 2020; Pasztoi & Ohnmacht, 2022) as well as *Bmp3, Efnb2*, and *Procr* (encoding Cd201), previously reported as a CTF marker in the colon (Melissari et al., 2021). We named the cluster 4 cells Sox6-CTFs (Figure 1C-D). Finally, cluster 5, in addition to the CTF-like expression pattern, specifically expressed *Il1rl1* (encoding for ST2, the IL-33 receptor) along with inflammation and tissue growth-associated genes such as *Ptgs2*, *Fgf7*, *Il11,* and *Stc1*. We named the cluster 5 ST2-CTFs (Figure 1C-D).

We have previously shown that Interleukin-11 (IL-11) expression is increased in polyps of PJS patients and mouse models (Ollila et al., 2018), and IL-11 is known to induce *Il1rl1* expression in fibroblasts (Widjaja, Chothani, et al., 2022). Moreover, increased expression of COX2, encoded by *Ptgs2,* has also been associated with PJS tumorigenesis (Udd et al., 2004), and stromal PTGS2 was previously shown to promote intestinal tumorigenesis (Roulis et al., 2020), suggesting functional importance for the ST2-CTFs. Indeed, the ST2^+^ fibroblast subset was almost exclusively found in polyps (Figure 1E). Interestingly, also the Wnt4-CTF population was enriched in polyps and expressed low levels of the ST2-markers *Il11* and *Il1rl1* in the polyp context, suggesting that ST2^+^ fibroblasts may represent an activated state of the Wnt4-CTFs (Figure 1F). The Pi16-CBF population, on the other hand, was primarily derived from control tissue (Figure 1E), consistently with the likely localization of the Pi16^+^ universal fibroblasts in the submucosa. Tissue staining confirmed high and mesenchyme-specific expression of the ST2 protein in polyps of *Twist2-Cre;Lkb1^fl/+^* as well as the *Lkb1^+/-^* mice (Figure 1G). The staining was most prevalent in the stroma of polyp edges, verifying the crypt top identity of the ST2-CTFs (Figure 1G).

In summary, we found that the fibroblast subsets in antral mucosa can be classified similarly to previously characterized intestinal and colonic fibroblasts but with specific differences, such as the lack of specificity in the *Cd81* expression (Table 1). Notably, one of the fibroblast clusters, the ST2-CTF population, was significantly enriched in tumors, suggesting these cells are tumor-promoting cells in PJS.

**Table 1.**
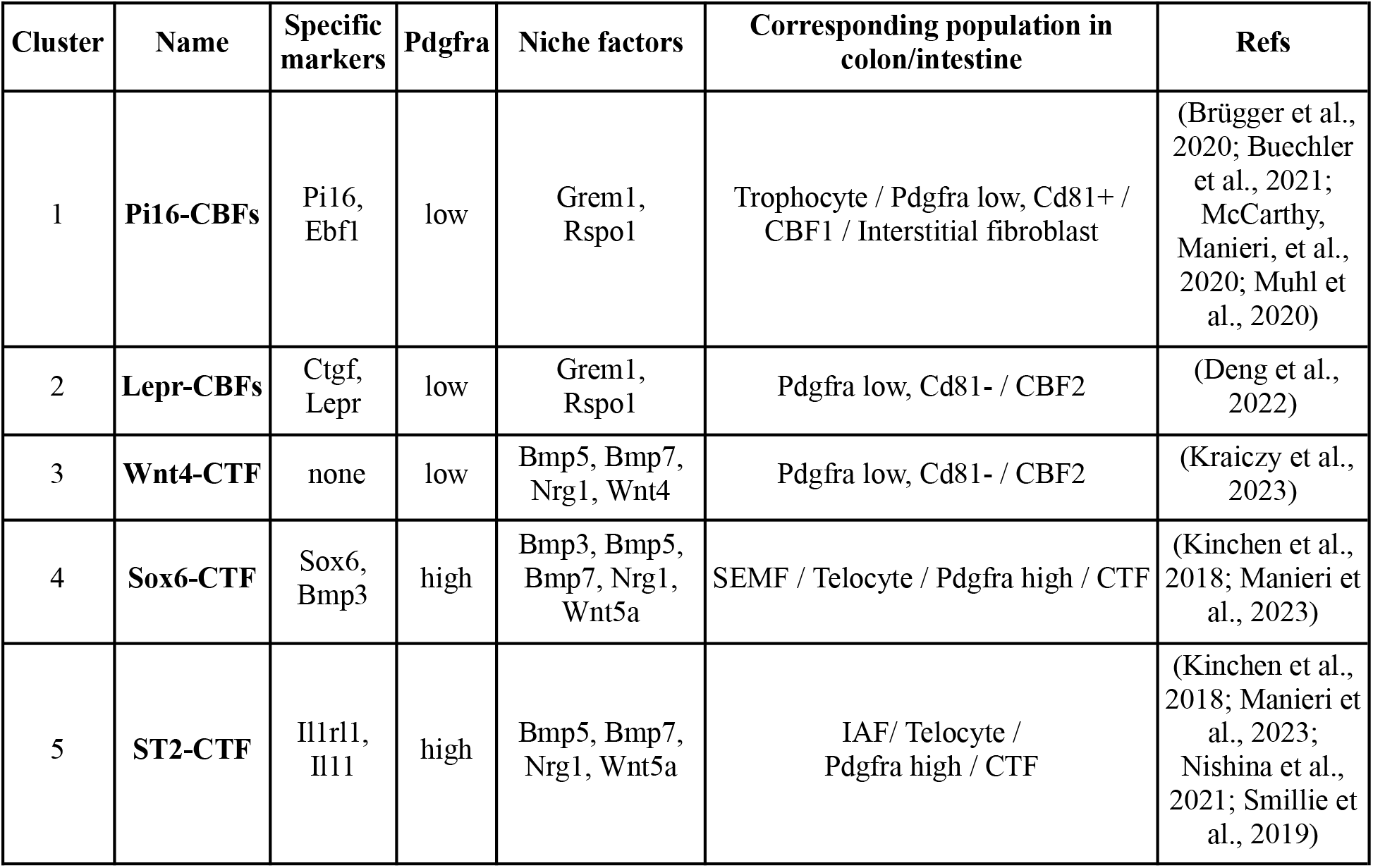
Identified gastric fibroblast subpopulations, top markers, and corresponding populations in the colon and intestine.

## CTF-specific loss of LKB1 induces polyposis and early expression of *Il1rl1* and *Il11*

Based on our finding that ST2-CTFs were enriched in PJS tumors, we aimed to investigate whether depleting Lkb1 in CTFs was sufficient to induce ST2-CTF genes and polyp formation. To address this, we crossed the *Lkb1^fl^*mice with *Foxl1-Cre* mice that restrict the Cre activity to subepithelial fibroblasts, representing the CTFs (Aoki et al., 2016) (Figure 1C). The *Foxl1-Cre;Lkb1^fl/fl^* mice started showing signs of discomfort at 4 months of age, and autopsies confirmed extensive gastric polyposis that severely obstructed the GI tract (Figure 2A-B). Interestingly, mice with heterozygous loss of Lkb1 (*Lkb1^+/-^* or *Twist2-Cre;Lkb1^fl/+^*) develop polyps with significantly slower kinetics and survive until about one year of age (Ollila et al., 2018; Rossi et al., 2002). The fast polyposis in *Foxl1-Cre;Lkb1^fl/fl^*mice confirmed our previous finding that homozygous loss of *Lkb1* in fibroblasts dramatically accelerates tumor formation (Ollila et al., 2018). Importantly, lineage tracing demonstrated that stromal loss of *Lkb1* in *Foxl1*-expressing cells resulted in an expansion of *Lkb1-*deficient fibroblasts in the polyp stroma (Figure 2C).

**Figure 2.**
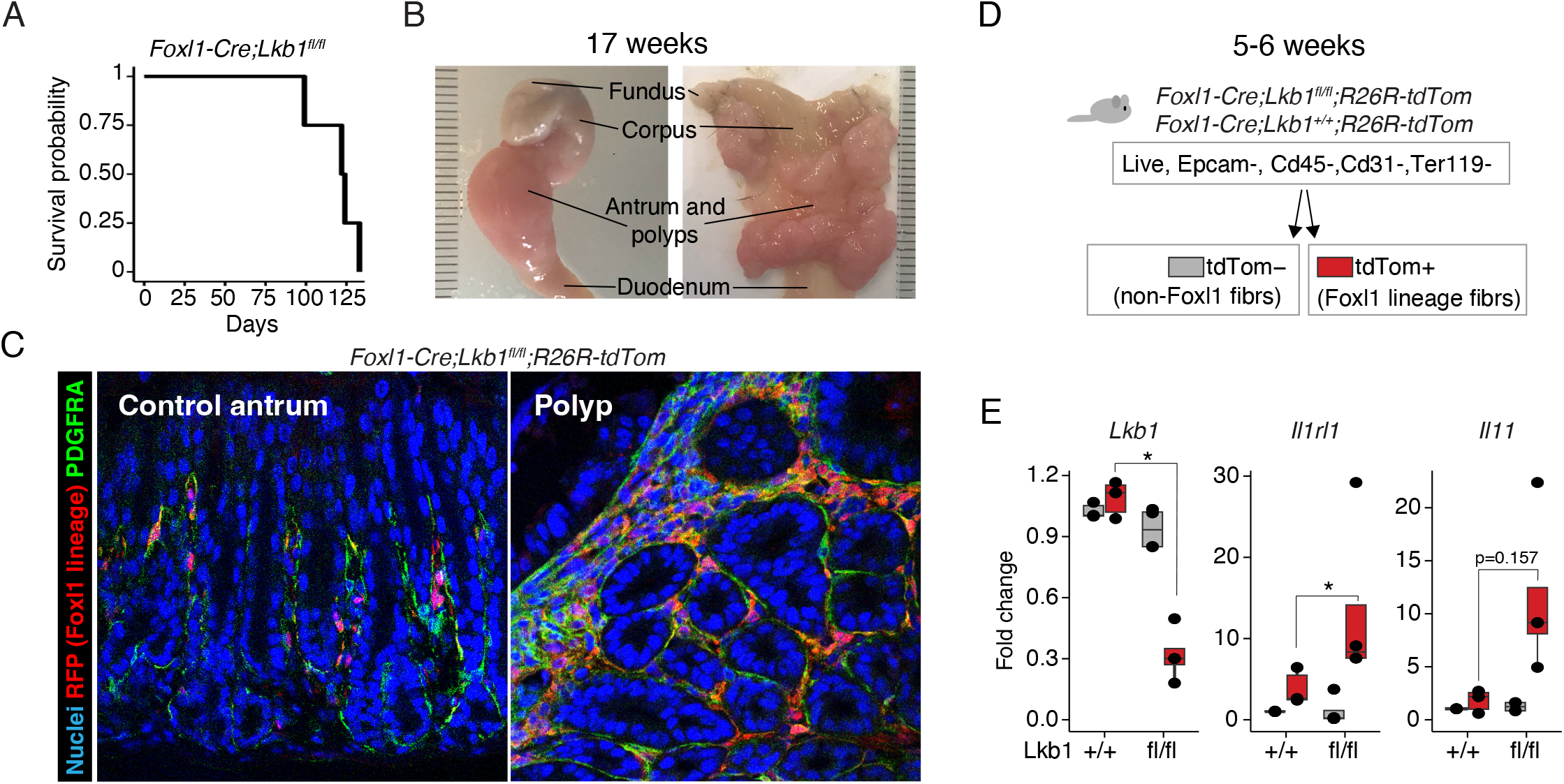
CTF-specific Loss of LKB1 induces polyposis and expression of ST2-CTF markers. **(A)** Survival curve of the Foxl1-Cre;Lkb1fl/fl mice (n=4). **(B)** Representative images of a stomach from *Foxl1-Cre;Lkb1fl/fl* mouse at 17 weeks. **(C)** Representative histology images of tumor and adjacent mucosa stained for PDGFRA (Fibroblasts) and RFP (Foxl1 lineage) from *Foxl1-Cre;Lkb1fl/fl;R26R-TdTomato* mouse at 17 weeks of age. **(D)** FACS strategy to isolate Foxl1-lineage fibroblasts from young mice of indicated genotypes. **(E)** Expression of Lkb1, Il1rl1 and Il11 mRNA in the indicated fibroblast fractions. *, p>0.05, one-tailed T-test.

To investigate if LKB1 loss induced the expression of the polyp-enriched ST2-CTF marker genes independently of polyp formation, we sorted gastric fibroblasts from young healthy *Foxl1-Cre;Lkb1^fl/fl^;R26R-tdTomato* and control mice prior to tumor formation. Loss of *Lkb1* expression and concomitant increase of the ST2 markers *Il11* and *Il1rl1* was observed in the *Foxl1-* lineage fibroblasts of the *Lkb1^fl/fl^* mice, showing that the ST2-CTF-like signature is a primary result of LKB1 loss (Figure 2D-E).

## LKB1 loss recapitulates the inflammatory ST2-CTF transcriptome

The ST2-CTFs expressed inflammatory and repair-associated genes (Figure 1D), and we sought to compare the ST2-CTF signature (top 25 genes expressed in the ST2-CTFs, Supplementary Table 1) with publicly available datasets related to GI injury and inflammation. The ST2 signature was strongly induced in the scRNAseq analysis of mouse colon fibroblasts treated with dextran sodium sulfate (DSS), inducing acute colon inflammation (Kinchen et al., 2018) (Figure 3A). Significant enrichment was also observed when the ST2 signature was compared to lipopolysaccharide (LPS)-treated primary intestinal fibroblasts, as well as with DSS-treated colon tissue lysates (Figure 3B). The ST2 signature was also enriched in inflammation-associated fibroblasts of ulcerative colitis patients (Smillie et al., 2019) (Figure 3C-D), indicating the signature is induced in inflammatory/regenerative fibroblasts in both mouse and human GI tract.

**Figure 3.**
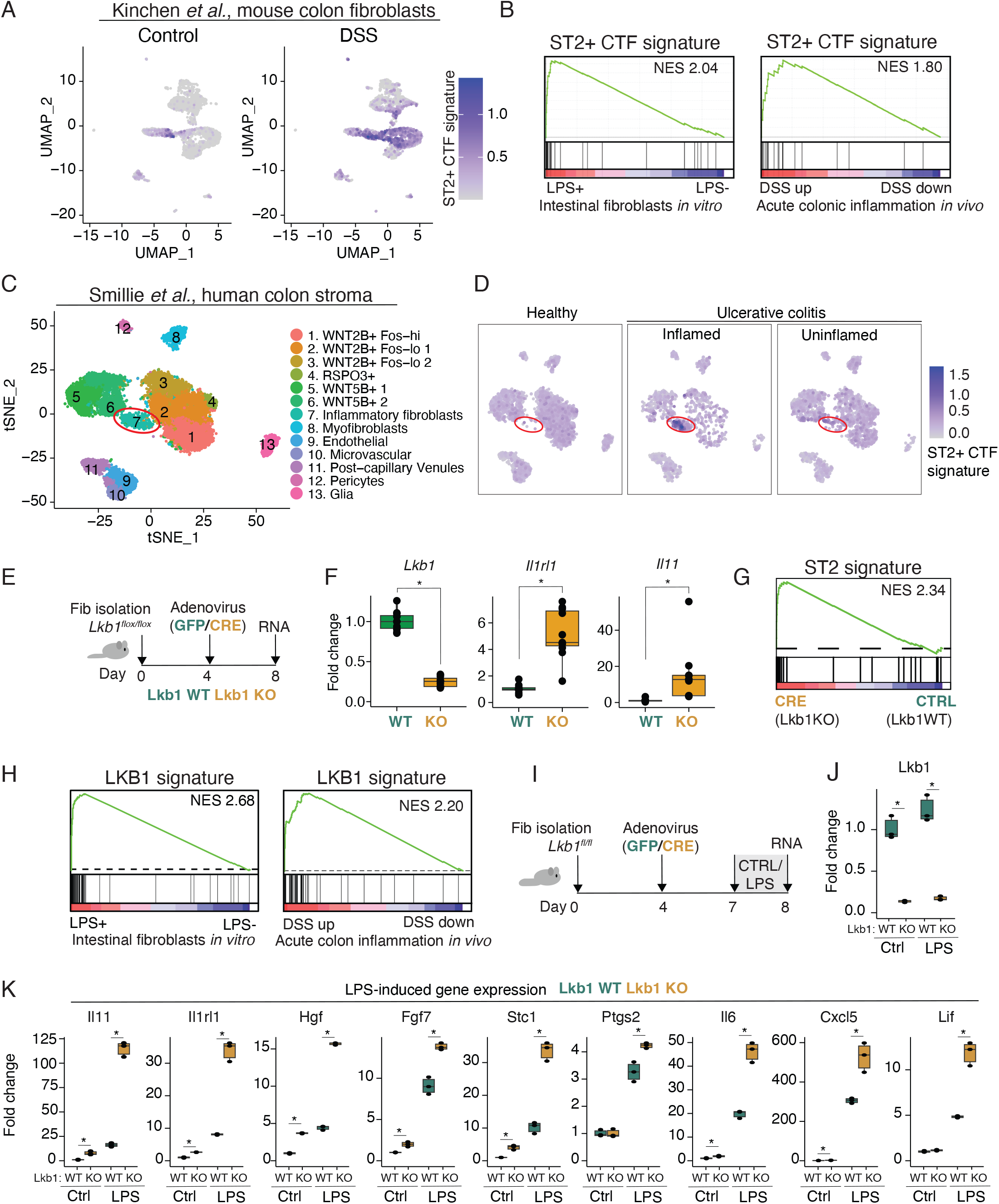
LKB1 loss recapitulates the inflammatory ST2-CTF transcriptome. **(A)** Feature plot of the ST2+ CTF gene signature in homeostatic (Control) and inflamed (DSS) mouse colon fibroblasts. Data from(Kinchen et al., 2018). **(B)** GSEA of ST2+ CTF gene set against LPS treated primary fibroblasts and DSS treated mouse colon tissue lysates. **(C-D)**. UMAP **(C)** and feature plot **(D)** showing the ST2+ CTF gene signature in human colon stroma from ulcerative colitis patients ((Smillie et al., 2019). **(E)** Outline of the acute *in vitro* deletion of *Lkb1* using adenovirus in primary intestinal fibroblasts derived from *Lkb1^fl/fl^* mice. **(F)** mRNA expression of *Lkb1, Il1rl1 and Il11* primary intestinal fibroblasts treated with adenovirus (Ctrl, Adeno-GFP; Cre, Adeno-Cre). Data from four independent experiments with two technical replicates each is shown. Average of the control replicates per experiment is used for normalization. **(G)** GSEA of the ST2+ gene signature against the ranked transcriptome from *Lkb1^fl/fl^* primary intestinal fibroblasts. **(H)** GSEA of LKB1 loss induced gene signature against LPS treated primary fibroblasts and DSS treated mouse colon tissue lysates. **(I)** Experimental setup and for assessing LPS-induced gene expression differences in WT and Lkb1 KO primary intestinal fibroblasts. Lkb1 mRNA levels as compared to Adeno-GFP in control conditions are shown. **(J)** mRNA expression of *Lkb1* in control and LPS-treated primary intestinal fibroblasts transduced with adenovirus (Ctrl, Adeno-GFP; Cre, Adeno-Cre). Data from four independent experiments with two technical replicates each is shown. **(K)** Expression levels of indicated genes and genotypes in control and LPS induced conditions. Fold change to WT Control cells is plotted. A representative of three independent experiments is shown. *, p<0.05, Two-tailed T-test. Box plots indicate mean and standard deviation.

Consistently with our observations linking the ST2^+^ fibroblasts to inflammatory fibroblasts, it was recently reported that *Lkb1* loss sensitizes mouse embryonic fibroblasts (MEFs) to inflammatory stimuli, such as IL-1β and LPS, resulting in enhanced induction of cytokine and chemokine expression (Compton et al., 2023). Thus, we investigated the potential role of inflammation in *Lkb1-*deficient fibroblasts and PJS tumorigenesis. In PJS, hamartomatous polyps develop throughout the GI tract, indicating that the underlying mechanisms are shared, and small intestinal polyps also develop in *Lkb1^+/-^* mice (Bardeesy et al., 2002) and *Twist2-Cre;Lkb1^fl/+^*mice (our unpublished observations). We cultured adult primary intestinal fibroblasts from *Lkb1^fl/fl^* mice and deleted *Lkb1* using adenoviral delivery of CRE recombinase, while control cells were transduced with a green fluorescent protein (eGFP)-encoding virus (Figure 3E). The transcriptional effects were assessed by quantitative PCR and bulk RNA sequencing. As expected, *Lkb1*-deficiency resulted in robust up-regulation of *Il1rl1* and *Il11* (Figure 3F), and the transcriptome was highly enriched for the ST2-CTF signature (Figure 3G). Similarly to the ST2-CTF signature (Figure 3B-C), the *Lkb1*-deficient fibroblast signature (Supplementary Table 1) was significantly enriched for inflammation-associated datasets, indicating that *Lkb1* deletion is sufficient to recapitulate the transcriptional features of inflammatory/regenerative fibroblasts (Figure 3H), consistently with the increased *Il11* and *Il1rl1* expression in *Lkb1*-deficient fibroblasts of tumor-free mice (Figure 2D-E). Next, we studied the inflammatory response of *Lkb1*-deficient intestinal fibroblasts by exposing them to LPS (Figure 3I-J). Interestingly, LPS treatment resulted in robust further induction of *Il11* and *Il1rl1* as well as other ST2 signature genes (*Hgf*, *Fgf7*, *Sct1* and *Ptgs2*), IL6-family cytokines (*Il6*, *Lif*) and the chemokine *Cxcl5* (Figure 3K), indicating that the *Lkb1* loss induced inflammatory phenotype can be further stimulated by exogenous inflammatory factors in primary intestinal fibroblast, similarly to previously shown in MEFs (Compton et al., 2023), but with a more robust response for *Il11* as compared to *Il6*. Of note, the expression of most investigated genes was significantly increased in *Lkb1*-deficient cells already without LPS treatment (Figure 3K).

## Inflammation exacerbates the LKB1 loss-mediated tumor-promoting phenotype

The higher expression of the LPS response genes in *Lkb1*-deficient cells suggested a functional role for inflammation in promoting PJS polyposis. To address if the inflammatory phenotype of fibroblasts has an effect on epithelial growth, we set up an organoid-fibroblast co-culture assay where wild-type (WT) or *Lkb1*-deficient primary intestinal fibroblasts were embedded into a 3D matrix (Matrigel) with WT intestinal epithelial organoids (Figure 4A-B). Intestinal epithelial organoids grow in Matrigel as three-dimensional structures recapitulating the homeostatic crypt-villus architecture of intestinal epithelium (Sato et al., 2009), but tumorigenic and regenerative organoids fail to form the correct 3D-structure and instead grow as hyperproliferative spheroids (Langlands et al., 2018; Nusse et al., 2018). As determined by eye and quantified using our automated organoid classification tool “Tellu” (Domènech-Moreno et al., 2023), *Lkb1*-deficient fibroblasts induced an increased frequency of spheroids as compared to organoids in the cocultures, indicating that a regenerative/tumorigenic phenotype of the WT intestinal epithelium is promoted by *Lkb1*-deficient fibroblasts (Figure 4C). This was consistent with *Lkb1*-deficient fibroblasts promoting polyp formation *in vivo*. The spheroid formation was further induced by LPS treatment in both WT and *Lkb1*-deficient fibroblast cocultures, suggesting that inflammation-induced growth-promoting signals from fibroblasts to epithelium are recapitulated by LKB1 loss and further boosted by inflammatory stimuli (Figure 4A-C).

**Figure 4.**
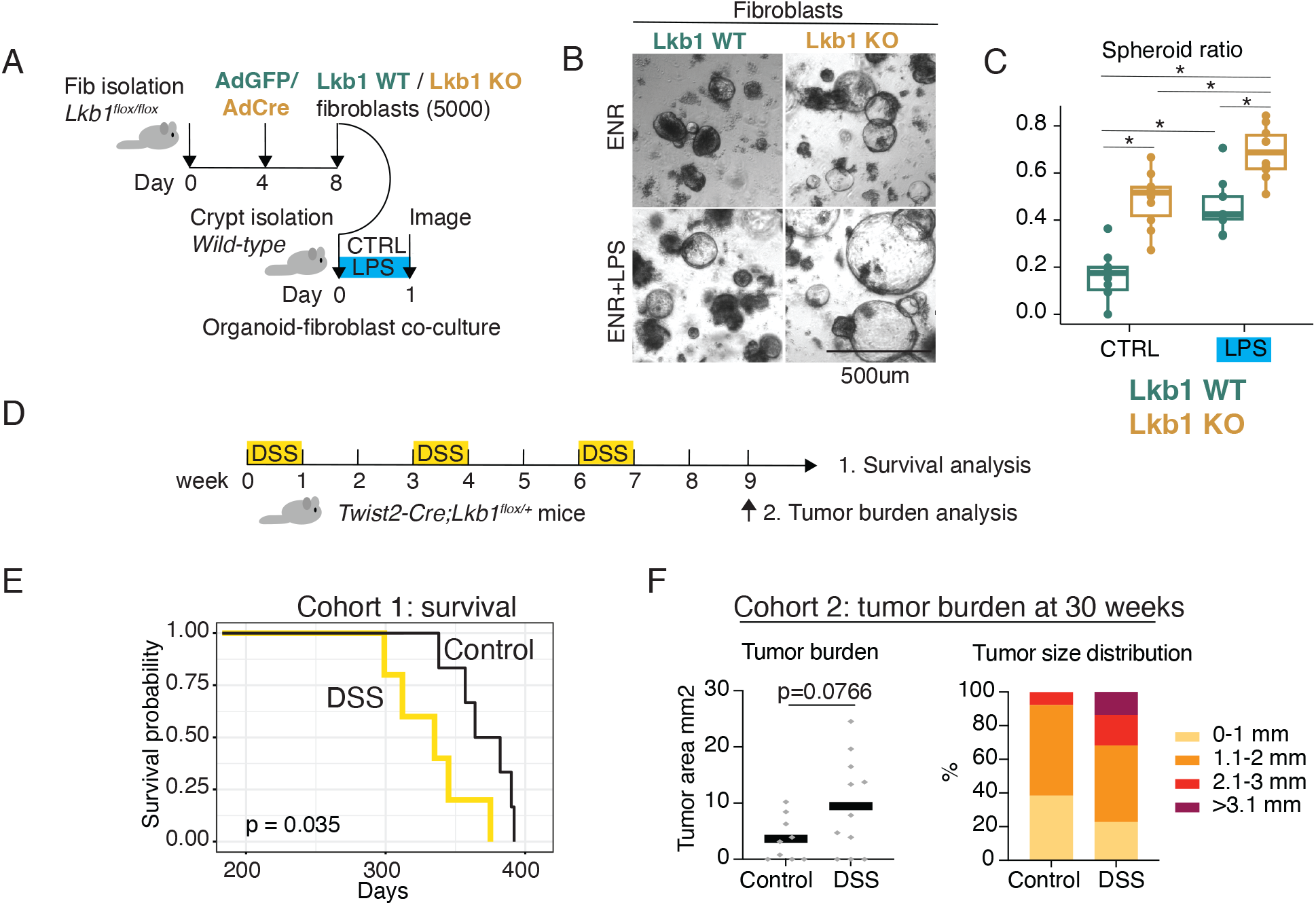
Inflammation exacerbates the LKB1 loss mediated tumor-promoting phenotype. **(A)** Experimental setup of the in vitro co-culture experiments using primary fibroblasts from *Lkb1fl/fl* mice and organoids isolated from wild-type mice. **(B)** Representative images of organoids after 24h of co-culture with Lkb1 WT (AdGFP) or Lkb1 KO (AdCre) fibroblasts with or without LPS. **(C)** Quantification of the ratio of spheroids vs organoids per well in each condition. The mean relative area per well, standard deviation and individual data points are shown (a.u, arbitrary units). **(D)** Experimental outline of the chronic inflammation induced in *Twist2-Cre;Lkb1fl/+* mice. **(E)** Kaplan-Meier curve depicting survival of *Twist2-Cre;Lkb1fl/+* mice in control conditions (black line, n=5) and upon DSS treatment (yellow line, n=6) Log-rank test used. **(F)** Tumor burden (left) and tumor size distribution (right) of the *Twist2-Cre;Lkb1fl/+* mice in control conditions (control, n=9) and upon DSS treatment (n=11).

To address the potential impact of inflammation-induced fibroblast reprogramming in PJS polyposis *in vivo*, we subjected *Twist2-Cre;Lkb1^fl/+^* mice to DSS-induced chronic inflammation (Figure 4D). Notably, while DSS is best known to result in colonic inflammation, the entire GI tract, including the stomach, is affected (Elsheikh et al., 2012). We observed a significantly reduced survival of DSS-treated *Twist2-Cre;Lkb1^fl/+^*mice (Figure 4E), linked with an increased gastric tumor size and a trend towards increased tumor burden (Figure 4F). Furthermore, subjecting the mice to a carcinogen (Azoxymethane, AOM) followed by DSS inflammation led to significantly increased gastric tumor burden in *Twist2-Cre;Lkb1^fl/+^*mice (Supplementary Figure 2A-B). The colonic tumor burden induced by AOM-DSS was also more prominent in *Twist2-Cre;Lkb1^fl/+^*mice compared to WT controls, albeit the difference did not reach statistical significance (Supplementary Figure 2C). Altogether, our results indicate that while exogenous inflammation is not required for PJS polyposis, it boosts the tumorigenic potential of LKB1 deficient mesenchyme, suggesting inflammation as a contributing mechanism driving polyp development.

## ST2-CTFs present a unique signaling pattern targeting fibroblasts and epithelial cells

Next, we set out to take a broader look into our scRNAseq data and also included the non-fibroblast cell populations. Epithelial (*Epcam^+^*) cells included surface cells (*Muc5ac^+^*), proliferative isthmus cells (*Mki67^+^*), gland and base cells (*Aqp5*^+^, *Muc6*^+^), as well as tuft cells (*Dclk1*^+^, *Trpm5*^+^), identified using previously established markers (Haber et al., 2017; Tan et al., 2020; Xiao & Zhou, 2020) (Supplementary Figure 3A-B). The distribution of the cell types was similar between normal antrum and polyp, in agreement with hamartomatous polyp type (Supplementary Figure 3C). However, the differential expression analysis of the epithelial cell fraction revealed some differences, such as upregulation of the regenerative markers *Reg3b* and *Reg3g* in polyps, confirming the epithelial origin of these changes noted in previous analyses of polyps lysates (Ollila et al., 2018) (Supplementary Figure 3D). In addition, we observed significantly reduced epithelial expression of *Pgc* (encoding for Pepsinogen C) and *Muc5ac* (encoding for the secreted Mucin 5AC) in the polyps, indicating that some of the previously observed Lkb1 loss-driven epithelial differentiation defects (Udd et al., 2010) are at least in part driven by stromal cells (Supplementary Figure 3D). Reduced expression of Pepsinogen C was confirmed using immunohistochemistry (Supplementary Figure 3E). Analysis of immune cells (Supplementary Figure 4A-C) and endothelial cells (Supplementary Figure 4D-E) identified all major cell types (James et al., 2022; Kalucka et al., 2020) and suggested a putative increase of neutrophils in the polyp-derived clusters as previously reported by (Poffenberger et al., 2018) (Supplementary Figure 4C, D).

Next, we analyzed cell-cell communication across all cell clusters identified in the control and polyp datasets using CellChat (Jin et al., 2021). This analysis revealed increased interactions within the polyp compared to the control antrum, suggesting activated signaling (Supplementary Figure 5A). Next, we plotted the total signaling strength from all cell types in a 2D space. Interestingly, the strongest outgoing and incoming signaling was observed in fibroblasts and was highest in the CTF clusters, including ST2-CTFs, suggesting that CTFs represent the most active cell types regarding paracrine signaling in our dataset (Figure 5A). We then further analyzed the outgoing and incoming cell-cell communication patterns across clusters. Five outgoing communication patterns were identified, mostly clustering along the major cell types (epithelial cells, endothelial cells, myeloid and lymphoid lineage immune cells, and fibroblasts) (Figure 5B). Interestingly, the tumorigenic ST2^+^ fibroblasts exhibited a unique outgoing communication pattern, Pattern 5, separating from the signaling pattern of other fibroblast subsets that signaled according to Pattern 1. The main differences were the active IL6 family signaling and parathyroid hormone (PTH) family signaling, as well as increased signaling from NRG, HGF, and FGF families, suggesting that emergence of the ST2-CTFs cells induces an altered stromal niche (Figure 5B). Interestingly, the only ligand contributing to the IL-6 family signaling was IL-11 (Supplementary Figure 5B). Upon closer examination, we identified distinct receivers of the ST2-CTF outgoing cell communication. The signals to epithelial cells included NRG and HGF family ligands, while PTH was received by pericytes and IL-11 signal by fibroblasts (Figure 5B-C). Interestingly, analysis of the ligand-receptor pairs constituting the signaling patterns predicted HGF-MET and NRG1-ERBB3 as the most active pathways emanating from ST2-CTFs to epithelial cells, suggesting functional relevance as noted before in regenerative and tumor contexts (Jardé et al., 2020; Joosten et al., 2017; Lemmetyinen et al., 2023) (Supplementary Figure 5B-C). FGF7-FGFR1 signaling was predicted to mainly target fibroblasts, while EREG-EGFR signaling was active to both epithelial cells and fibroblasts. Interestingly, the activation of IL-6 family signaling was attributed to increased expression of IL-11 signaling from ST2-CTFs to IL11RA1-IL6ST (Gp130) receptor-expressing fibroblasts populations (Supplementary Figure 5B-D). This observation suggested an upstream role for IL-11 in fibroblast reprogramming.

**Figure 5:**
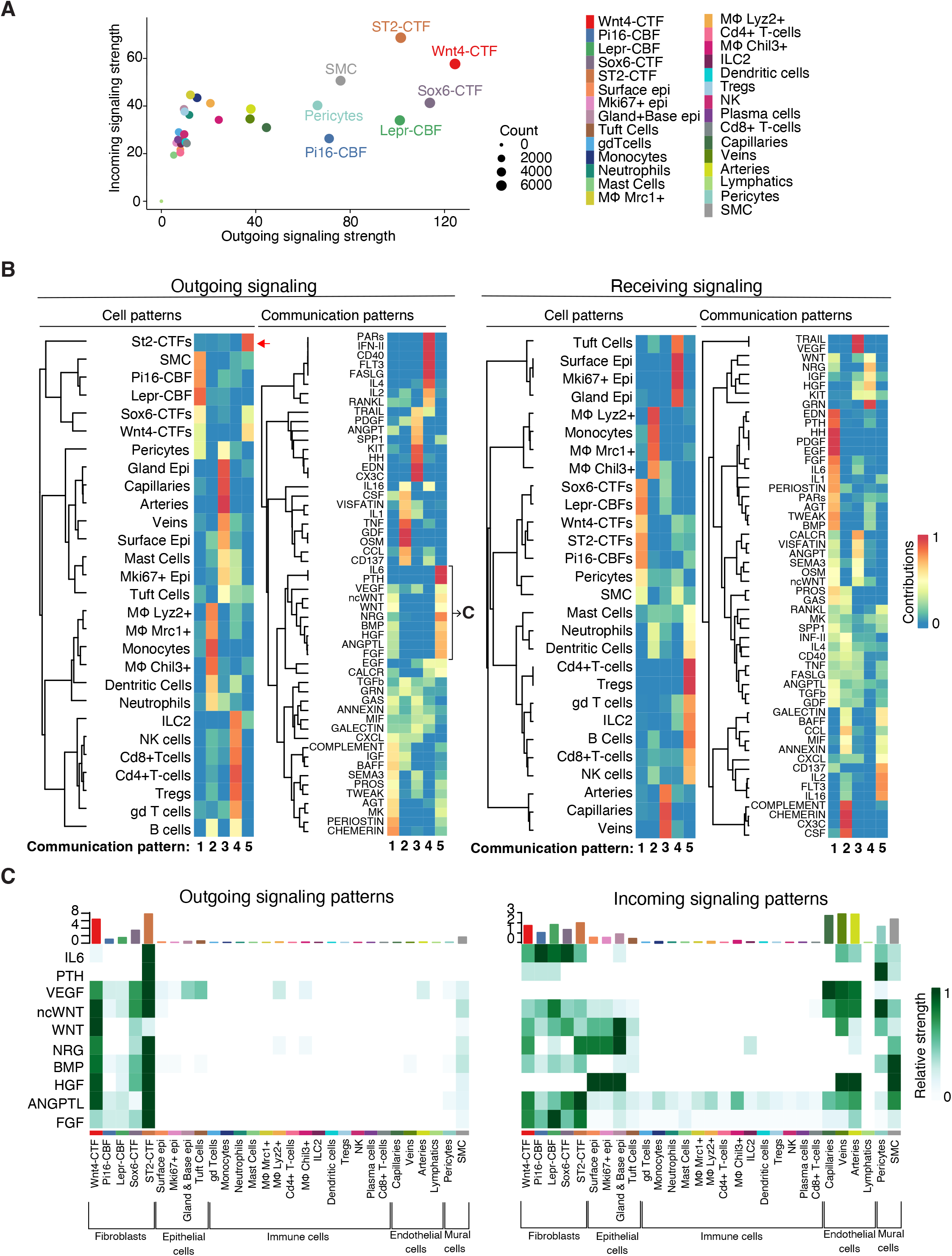
ST2-CTFs possess a strong and unique cell-cell signaling pattern. **(A)** 2D representation of overall incoming and outgoing interactions in the tumor gastric mucosa. **(B).** Heatmap of the cell communication pattern analysis of outgoing and receiving signals between clusters. **(C)** Heatmap of the cell communication outgoing signal pattern 5 found in B specific of ST2^+^ fibroblast.

## Inhibition of IL-11 blocks the ST2^+^ signature and reduces polyposis

The scRNAseq data suggested a role for IL-11 in fibroblast reprogramming (Figure 5C, Figure 6A). Consistently, the *Lkb1*-deficient intestinal fibroblast signature was enriched in the transcriptome of IL-11-expressing fibroblasts isolated from inflammation-induced mouse colon tumors (Nishina et al., 2021), indicating shared functional outcomes of IL-11 expression and LKB1 loss (Figure 6B). To explore whether the *Lkb1*-deficient ST2-CTF-like fibroblast transcriptome was dependent on IL-11, we engineered *Il11* knock-out (KO) immortalized intestinal fibroblasts generated from *Lkb1^fl/fl^* mice using CRISPR-Cas9 (Supplementary Figure 6A). Remarkably, the induction of the ST2^+^ signature observed upon Cre-mediated *Lkb1* loss was significantly reduced in the absence of *Il11* (Figure 6C, Supplementary Figure 6B), implying that the *Lkb1*-loss mediated reprogramming of intestinal fibroblasts into the ST2-CTF-like phenotype is at least partially dependent on IL-11.

**Figure 6.**
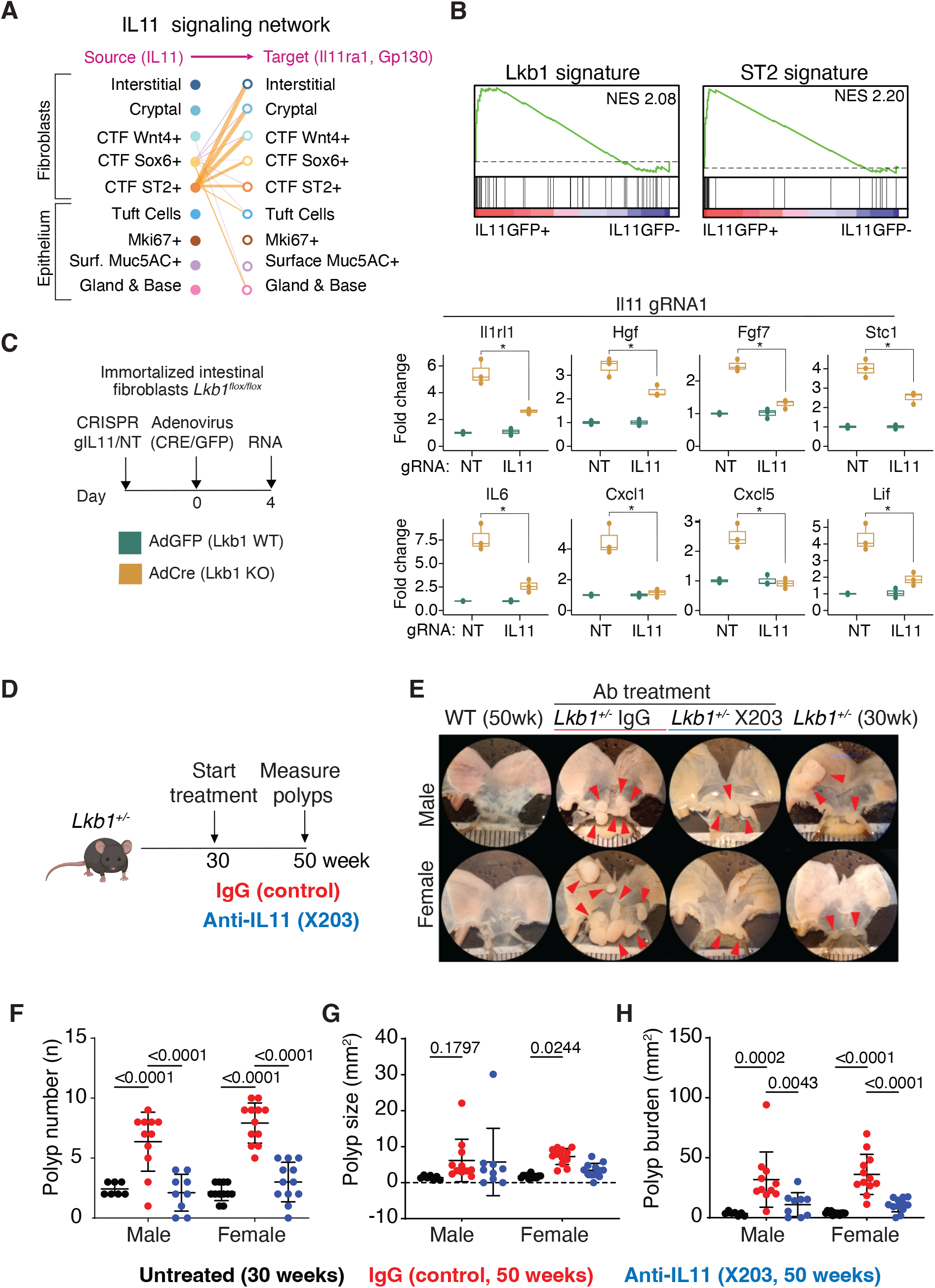
Inhibition of IL-11 signaling inhibits the ST2+ transcriptome and reduces PJS polyposis. **(A)** IL11 cell-cell signaling network originates from ST2-CTFs and targets fibroblast populations in PJS tumors. **(B)** GSEA of Lkb1 and ST2+ fibroblast gene sets (Supplementary Table 1) against the transcriptome of IL11-EGFP+ fibroblasts isolated from inflammation-associated colon cancer mouse model (Nishina et al., 2021). **(C)** Experimental setup (left) and gene expression of indicated genes in fibroblasts transduced with Cas9 and non-targeting (NT) gRNA or gRNA against IL11 (right). gRNA1 is shown, see also Supplementary Figure 6. **(D)** Schematic diagram illustrating antibody treatment in *Lkb1*^+/-^ mice. 30-week-old *Lkb1*^+/-^ mice with established polyposis were treated with either X203 or IgG isotype antibodies (20 mg/kg/week IP) up to 50 weeks (n=11 males, 12 females per antibody group). **(E)** Representative images of stomachs of 50-week-old *Lkb1*^+/-^ mice treated with either antibody compared to 50-week-old wildtype littermates and 30-week-old untreated *Lkb1*^+/-^ mice. Red arrows indicate gastric polyps. **(F-H)** Average polyp number **(F)**, size **(G)**, and total polyp burden **(H)** of *Lkb1*^+/-^ mice treated with either antibody compared to untreated 30-week-old *Lkb1*^+/-^ mice (n=7 males, 12 females).*, p<0.05, **, p<0.01, ***, p<0.001, ****, p<0.0001, two-way ANOVA with Sidak post-hoc test. ns, not significant.

Interestingly, previous work has indicated a crucial role for IL-11 in cardiorenal fibrosis (Schafer et al., 2017), and a blocking antibody to IL-11, X203, inhibits fibroblast activation *in vitro* and *in vivo* (Ng et al., 2019). We hypothesized that blocking IL-11 in mice modeling PJS could effectively inhibit fibroblast reprogramming and tumorigenesis following *Lkb1* inactivation. To address this, the *Lkb1^+/-^* mice were treated with X203 or IgG intraperitoneally at 20 mg/kg dose weekly from 30 to 50 weeks of age (Figure 6D). Notably, X203 treatment significantly reduced the polyp number and polyp burden while not affecting individual polyp size, implicating that IL-11 signaling plays a significant role in PJS polyp initiation (Figure 6 E-H). These results indicate that the LKB1 loss-mediated fibroblast reprogramming leading to PJS tumorigenesis is driven by therapeutically targetable expression of IL-11.

## Discussion

Here, we investigated fibroblast heterogeneity and cell-cell interactions driving PJS polyposis using mouse models. Interestingly, the polyps were enriched for a tumorigenic fibroblast population, ST2-CTFs, presenting with a high expression of growth factors and inflammatory-associated genes such as *Il11*, *Il1rl1, Fgf7, Hgf*, *Stc1*, and *Ptgs2*. Importantly, loss of LKB1 was sufficient to induce an ST2-type inflammatory phenotype in the fibroblasts, and inflammatory insults further enhanced this phenotype. Thus, while LKB1 loss alone altered the fibroblast function, further activation via local inflammation may be involved in the etiology of PJS polyp development, also supported by the recent report on *Lkb1*-deficient MEFs being sensitized to inflammatory stimuli (Compton et al., 2023) and other reports connecting LKB1 loss to increased inflammation (Ollila et al., 2018; Poffenberger et al., 2018). Interestingly, the ST2-CTFs represented only a subset of fibroblasts in the polyps, densely populating the polyp edges. Since all fibroblasts in the polyps were heterozygous for LKB1 in our model, this observation suggests that additional insults, perhaps originating from GI lumen, are required for the fully activated ST2-CTF phenotype. This poses an interesting question of whether mechanisms besides genetic LKB1 loss, such as inhibitory phosphorylation (Widjaja, Viswanathan, et al., 2022), is associated with fibroblast activation in inflammatory conditions. Since PJS patients are heterozygous for LKB1, and the mouse models with homozygous stromal *Lkb1* loss present with faster polyp formation (Ollila et al., 2018), this study), inhibition of the wild-type LKB1 allele could contribute to further fibroblast activation also in PJS polyposis. This would be an interesting topic for a future study.

GI fibroblast heterogeneity has been under intensive investigation in recent years (Brügger et al., 2020; Deng et al., 2022; Jasso et al., 2022; Kinchen et al., 2018; Kraiczy et al., 2023; Manieri et al., 2023; McCarthy, Manieri, et al., 2020; Pærregaard et al., 2023) We found that antral fibroblasts clustered similarly to intestinal and colonic fibroblasts, including the formation of Wnt and Bmp gradients (Brügger et al., 2020; McCarthy, Manieri, et al., 2020) but with differences in specific marker expression: for example, in the gastric mucosa *Cd81* was expressed across all fibroblast subsets, as opposed to intestinal and colonic fibroblasts where *Cd81* specifically marks one of the fibroblast population providing the trophic support for stem cells (Brügger et al., 2020; McCarthy, Manieri, et al., 2020). The corresponding population in our dataset was specifically marked by *Pi16*, also known as a marker of “universal fibroblasts’’ (Buechler et al., 2021; Muhl et al., 2020), consistent with submucosal location away from the tissue parenchyma. We also found a Lepr^+^ CBF population. Interestingly, the Lepr^+^ fibroblasts provide stem cell niche factors for hematopoietic stem cells (Comazzetto et al., 2019) and intestinal stem cells (Deng et al., 2022). Expression of *Grem1* and *Rspo3* in antral Lepr-CBFs suggested they represent the primary stem cell supporting fibroblasts also in the gastric antrum. Our analysis also revealed an intermediate fibroblast population, Wnt4-CTFs, with similarities to the recently published Wnt4^+^ fraction of the *Pdgfra*-low, *Cd81*^-^ cells in the colon and intestine (Kraiczy et al., 2023), suggesting that the division to *Grem1*-expressing and *Wnt4*-expressing *Pdgfra*-low mucosal fibroblasts is conserved along the GI tract. Cell-cell communication analysis predicted active BMP signaling between these fibroblast subsets (Figure 5C), suggesting that division to Lepr-CBFs and Wnt4-CTFs is regulated by BMP signaling also in the gastric antrum. The relevance of Wnt4 signaling to GI homeostasis is unclear, but previous reports suggest functions related to promoting differentiation (Eliazer et al., 2019; Zhang et al., 2021). The Sox6-CTF cluster identified in our analysis presented with markers suggesting a subepithelial location away from the crypt/gland bottom, consistent with previous data (Kinchen et al., 2018). Interestingly, SOX6 has been previously implicated in suppressing ϐ-catenin-mediated transcription, consistent with the localization away from the stem cell zone (Iguchi et al., 2007).

Our results indicate that Lkb1 loss-induced IL-11 signaling is critical for activating the ST2+ fibroblast phenotype. IL-11, a proinflammatory IL-6 family cytokine, is upregulated primarily in stromal fibroblasts in several non-malignant inflammatory diseases (Fung et al., 2022). Interestingly, persistent transgenic overexpression of IL-11 in mesenchymal cells was sufficient to elicit spontaneous GI inflammation leading to colonic fibrosis and mortality in mice, highlighting the relevance of careful regulation of IL-11 to avoid excess inflammation (Lim et al., 2020). In patients with ulcerative colitis, inflammatory-associated fibroblasts (IAF) characterized by high expression of IL-11 are expanded by more than 100-fold in inflamed tissue (Smillie et al., 2019), and IL-11-expressing fibroblasts emerge during inflammation-associated colon tumorigenesis in mice (Nishina et al., 2021). Interestingly, the IL11^+^ fibroblasts harbor a unique gene signature that predicts poor prognosis in human colorectal cancer (CRC) (Nishina et al., 2021), and activation of STAT3 through IL-6 or IL-11 in CRC cancer-associated fibroblasts (CAFs) is associated with poor prognosis (Heichler et al., 2020), suggesting relevance for IL-11 signaling in GI tumorigenesis beyond PJS. IL-11 was found to be significantly correlated with STAT3 activation (Putoczki et al., 2013) and thus, the observed therapeutic responses to JAK inhibitors in PJS models (Ollila et al., 2018; Poffenberger et al., 2018) could result from the inhibition of the JAK/STAT3-driven fibroblast activation. The JAK inhibition could also act on innate immune cells and reduce the production of inflammatory cytokines such as IL-1β, a potent activator of LKB1-deficient MEFs (Poffenberger et al., 2018).

Interestingly, our CellChat cell-cell communication analysis provided a detailed overview of the dynamic interplay between ST2-CTFs and the surrounding cellular components, including high IL-11 signaling to other fibroblast populations. Also, BMP, non-canonical WNT as well as FGF family signaling were predicted to affect other fibroblasts predominantly, and it would be interesting to address the contribution of these signals to the fibroblast activation, as well as to the spatial fibroblast distribution (Figure 5C, Supplementary Figure 5C). However, despite the critical role of fibroblasts, the PJS polyps consist mostly of highly proliferating epithelial cells (Ollila et al., 2018; Udd et al., 2010). Our finding that IL-11 signaling impacts the fibroblast phenotype raises the question of the mechanism driving the epithelial hyperproliferation. Interestingly, ST2-CTFs expressed a plethora of growth-promoting factors such as HGF, NRG1 as well as PTGS2 (the rate-limiting enzyme for Prostaglandin E2), known to promote the growth of GI epithelial cells (Ding et al., 2018; Jardé et al., 2020; Lemmetyinen et al., 2023; Roulis et al., 2020; Takahashi et al., 1995). Thus, ST2-CTFs appear to play a pivotal role in reprogramming the fibroblasts into an inflammatory/regenerative state, as well as driving the proliferative response observed in the epithelium, likely driven by a combination of growth-promoting factors. In addition, it remains possible that the upregulation of fibroblast-originating chemokines such as CXCL5, and the subsequent recruitment of immune cells such as neutrophils, plays a functional role in polyp formation. However, as the immune cells in our model are wild-type for LKB1, any putative polyp-promoting functions likely represent secondary changes.

Our discovery that IL-11 was required for the full inflammatory, ST2-fibroblast transcriptome induced by LKB1 loss points towards a feed-forward loop between LKB1 inhibition and IL-11 expression, along the lines of a previous report (Widjaja, Viswanathan, et al., 2022). Notably, the efficient blocking of polyp formation using a blocking antibody for IL-11 in PJS mouse models clearly indicates that IL-11 signaling is a key step in PJS tumorigenesis, which can be exploited therapeutically. As IL-11 expression is very low under normal homeostasis (Nishina et al., 2021), its targeting is expected to pose minimal side effects, opening a possible avenue for continuous treatment to reduce the need for surgical polyp removal, thus contributing to the quality of life in PJS patients. This calls for the initiation of clinical trials addressing the efficiency of IL-11 targeting in PJS.

In conclusion, by unraveling the interplay between IL-11 signaling and fibroblast activation following LKB1 loss, our study provides valuable insights into the tumorigenic mechanisms and therapeutic opportunities for PJS. Importantly, studying rare diseases often results in better understanding of common disorders, and therapies against rare conditions may also be useful in a wider context (Gahl, 2012). As the tumor microenvironment including cancer-associated fibroblasts are known to play an important role in sporadic GI cancer (Al-Bzour et al., 2023), it will be interesting to address whether our findings bear relevance also beyond PJS.

## Methods

### Animal experiments

The following mouse strains were used in this study: Twist2-Cre (The Jackson Laboratory #008712, (Yu et al., 2003)), Foxl1-Cre (The Jackson Laboratory #017738, (Sackett et al., 2007), Lkb1 flox (*Lkb1^fl^*) (The Jackson Laboratory #014143, (Bardeesy et al., 2002)), *Lkb1^+/-^* KO (Cyagen #S-KO-04594). Rosa26-tdTomato (The Jackson Laboratory #007914)(Madisen et al., 2010). Animals were housed at the University of Helsinki Laboratory Animal Center and the SingHealth Experimental Medicine Centre according to national and international legislation and guidelines and experiment approvals (ESAVI/3816/2023, 2019/SHS/1481 and 2019/SHS/1483). Mice had free access to food and water. Male and female mice with a similar gender ratio were used in all experiments unless otherwise stated.

### Antibody treatment in *Lkb1*^+/-^ mice

To evaluate putative beneficial effects of neutralizing IL-11 signaling, 30-week-old *Lkb1*^+/-^ mice (22 males and 24 females) were block randomized to receive either anti-IL11 (X203, Genovac) or an IgG isotype control (11E10, Genovac) at a dose of 20 mg/kg once per week IP. At 50 weeks, mice were euthanized by ketamine (100 mg/kg) and xylazine (10 mg/kg) mixture given IP, and the stomach extracted for polyp assessment. The stomach and proximal duodenum were opened, flushed with ice cold PBS and gently swapped with a cotton bud to remove debris, prior to image capture under the stereomicroscope (Olympus, SZ61). Gastric polyps were enumerated, and size quantified using ImageJ (version 1.53, NIH). Polyp size was reported as the average polyp area per stomach, and total polyp burden was assessed as the summation of individual polyps’ area per mouse. 30-week-old untreated *Lkb1*^+/-^ mice (7 males and 12 females) were included as a reference group prior to antibody treatment to gauge therapy effectiveness. For statistics, we performed a two-way ANOVA with Sidak post-hoc analysis for the two factors, treatment (30w untreated vs. 50w X203 vs. 50w IgG) and sex (males vs. females).

### GI inflammation and inflammation-induced tumorigenesis models

Male mice were used for the inflammation experiments. The acute GI inflammation was induced by replacing the drinking water of mice with 3% dextran sodium sulfate (DB001, TdBlabs) diluted in water for 7 days. The mice were euthanized after a 3-day recovery period with normal drinking water. The chronic inflammation was induced with three rounds of 7-day 3% DSS treatment and a 14-day recovery with normal drinking water. The mice’s weight was monitored daily throughout the experiment to monitor weight loss. The survival cohort of DSS-treated and control mice was followed, and the mice were euthanized when they started to show signs of discomfort (abnormal posture and reduced activity) and were autopsied to confirm large gastric polyps as the cause of death. The mouse cohort for tumor burden analysis was euthanized two weeks after the last DSS cycle. For the AOM-DSS inflammation-induced colorectal cancer model, mice were injected intraperitoneally with 10 mg/kg azoxymethane (AOM) in PBS, and after 7 days followed by three cycles of 7 days if 3% DSS and 14 days of recovery as described above. Gastric and colonic polyps were counted and measured under stereomicroscope (Leica MZFLIII).Gastric tissue dissociation and scRNAseq library preparation

Antral polyps and adjacent antral mucosa from a pair of mice were dissociated into single cells as follows: the tissue was flushed with PBS, minced into < 1 mm pieces with scissors, and placed in a 15 ml falcon tube filled with 5 ml of a collagenase mixed solution (collagenase II (Gibco 17101-019) + IV (ThermoFischer 17104019), 1:1 ratio at 2 mg/ml). The samples were incubated horizontally in a 37°C shaking incubator (220 rpm) for 40 min. During this incubation time, pieces were pipetted up and down 10 times every 10 min, and the collagenase solution was changed after 20 minutes, collecting the supernatant containing the single cells. The supernatant (∼10 ml) was then transferred to a clean Falcon tube, and collagenases were inactivated with cold DMEM supplemented with 20% fetal bovine serum (FBS). The samples were passed through a 70 um cell strainer, centrifuged for 10 min at 1100 rpm, and single cells were resuspended in 1–2 ml of PBS. Single-cell suspensions were stained with 7 AAD (LifeTechnologies #A1310) at a 1:1000 dilution prior to FACS sorting to exclude dead cells. 10.000 live cells per sample were sorted at ultra-purity using a Sony SH800Z Cell Sorter at the Biomedicum Flow Cytometry unit. The 10X v3 library kit (Library Construction Kit from 10X Genomics #1000190) was used for scRNA-seq library preparation. Samples were sequenced using a drop-seq-based 10X Genomics machine with a target of 5000 cells per sample.

### Isolation of primary intestinal fibroblasts

The small intestine was cleaned with cold PBS using a syringe and 21-gauge needle, cut longitudinally, and rewashed from intestinal contents with cold PBS. The tissue was then placed in a Petri dish and cut into 1 cm pieces with scissors and forceps, and tissue pieces were transferred into a 15 ml Falcon tube. To remove intestinal epithelial cells, 10 ml of pre-warmed EDTA solution (5 mM EDTA and 1 mM DTT) was added to the tissue and incubated for 20 min at 37°C in a shaking incubator with the tubes placed horizontally. The tissue pieces were washed four times with cold PBS by vigorously shaking the tubes for 15 sec and removing the supernatant each time. After the last wash, the supernatant was no longer cloudy. The tissue was further minced to <1 mm using scalpels or scissors. 10 ml of collagenase mix digestion solution (collagenase II + IV mixed at 1:1 ratio at 2 mg/ml -Thermo Fisher Scientific) was added to the tissue and incubated horizontally for 20 min at 37°C in a shaking incubator. The tissue was then pipetted up and down with a 10 ml pipette 15 times. 20 ml of cold fibroblast growth media (DMEM+20% FBS+1% penicillin/streptomycin+1% L-glutamine) was added to stop the reaction. The tissue was then centrifuged at 1400 rpm for 10 min, and the supernatant was removed. The pellet was resuspended in 10 ml of growth media and placed in a culture plate. The plate was washed three times with PBS after 1 h to remove floating cells and debris.

### Cell sorting

The antrum tissue was dissected and dissociated into single cells as described above (for scRNAseq tissue sample preparation). Single-cell suspensions were stained with 7-AAD at a 1:1000 dilution. Fibroblasts were enriched by negative selection excluding epithelial, endothelial, immune and red blood cells by staining the cell suspensions with antibodies against Epcam (eBioscience 11-5791-82), Pecam1 (eBioscience 11-0311-82), Cd45 (eBioscience 11-0451-82), and Tert19 (eBioscience 11-5921-82) conjugated to FITC at a 1:100 dilution for 30 minutes on ice. Sorting was performed using a Sony SH800Z Cell Sorter. Single cells were gated using forward and side scatter characteristics, and live cells were gated by excluding 7 AAD-positive cells. Fibroblasts were enriched by excluding the FITC-stained cells. The remaining cell suspension consisted mainly of fibroblasts, and fibroblasts from the Foxl1-Cre lineage were gated using the TdTomato channel. 10,000 TdTomato+ and TdTomato-cells of the fibroblast enriched fractions were sorted directly into Tri reagent® (TR118, Fisher Scientific), and RNA was extracted according to manufacturer’s instructions.

### Adenoviral transduction of primary intestinal fibroblasts

1.2 million primary intestinal fibroblasts were seeded in a 10 cm plate. The adenoviruses (AdCre (Iowa-5) or AdGFP (Iowa-4) at MOI 1000 were added to 6 mL of fibroblast growth media (see above), and the cells were incubated overnight with the virus. Following incubation, the media was removed, and the cells were washed once with the fibroblast growth media. The cells were used for experiments 4 days after the transduction.

### Co-culture of intestinal epithelial organoids and fibroblasts

Intestinal crypts were isolated from the small intestine of euthanized mice and cultured as previously described (Sato et al., 2009) with some modifications. Briefly, the small intestine was removed and mesentery was removed. The lumen was rinsed with ice-cold PBS using a syringe and gavage needle. The intestine was open longitudinally and gently rubbed to remove mucus. The small intestine was cut into 2-3 mm fragments with small scissors and the fragments were collected in a 50 ml Falcon tube and washed with PBS. The tube was then filled with 10 mM EDTA in PBS, and the tissue pieces were incubated for 45 minutes horizontally on ice. The EDTA-PBS solution was changed 3 times during this period, and gently shaking the tube each time. After an additional 60 minute incubation, the tube was shaken vigorously for 15 seconds to detach the crypts, and the contents were filtered through a 70 µm mesh into a fresh 50 ml Falcon tube to exclude villi and tissue pieces. The crypts were pelleted by centrifugation at 200 g for 2 min, washed with PBS and spun again to collect the crypts. The pellet was resuspended in ENR crypt media consisting of composed of basal organoid media (Advanced Dulbecco’s modified Eagle medium (DMEM):F12 (12634010, Gibco) supplemented with 1 M HEPES pH 7.4 (H0887, Sigma-Aldrich), 1× GlutaMAX (35050061, Gibco), 1× penicillin-streptomycin (P4333, Sigma-Aldrich), 1× N-2 supplement (17502001, Gibco) and 1× B-27 supplement (17504044, Gibco)) supplemented with EGF (Gibco, PMG8041; 50 ng/ml), NOGGIN (Peprotech, 250-38; 100 ng/ml), R-Spondin 1 (R&D Systems 3474-RS; 250 ng/ml) and N-acetylcysteine (A0737, Sigma-Aldrich; 1 µM). Y-27632 (Y0503, Sigma-Aldrich; 10 µM) was included in the ENR medium for the first two days. 20 µl domes containing 75% Matrigel (356231, Corning) and 25% crypt suspension were plated in pre-warmed 48-well plates. In total, approximately 200 crypts were plated per dome together. The domes were overlaid with 350 ul of ENR media and cultured at 37°C in a humidified atmosphere of 5% CO2. For co-cultures with fibroblasts, domes were overlaid with basal organoid media containing 5000 primary fibroblasts from *Lkb1^fl/fl^* mice previously adenotransducted with AdCre/AdGFP and incubated for 24 h with or without 1ug/ml LPS (Sigma L4516).

### Immunohistochemistry and immunofluorescence

Paraffin-embedded tissue sections were subjected to deparaffinization using a series of xylene and ethanol washes. Antigen retrieval was performed at 95°C for 25 minutes using Dako citrate buffer pH 9 (S2367) except for Pepsinogen C, where antigen retrieval was done with 0.05% Trypsin (215240, Difco) in 37C for 10 minutes, followed by 10 min in room temperature. The sections were washed with TBS and permeabilized with 0.1% Triton-X/PBS for 10 minutes at room temperature. A PAP pen was used to outline the sections, and blocking was done using 5% NGS in TBS for an hour at room temperature. Antibodies were incubated overnight at +4°C. Primary antibodies used were anti-RFP (1:500, 600-401-376, Rockland), anti-Pdgfra (1:100, AF1062, R&D Systems) and anti-St2 (1:150 AF1004, R&D Systems) and anti-Pepsinogen C. The following day, sections were washed with TBS and then incubated with a secondary antibody (Alexa Fluor-conjugated antibodies at 1:500) for one hour at room temperature. After washing again with TBS, counterstaining was performed using Hoechst (0.01ng/ml) for one minute, followed by rinsing with water and mounting in Mowiol. The stained sections were scanned using the Pannoramic 250 FLASH III digital slide scanner. Confocal imaging was performed with Leica TCS SP8 CARS Confocal microscope.

### Gene editing using CRISPR-Cas9

Mouse primary intestinal fibroblasts from passage 3 were transduced with a retrovirus encoding residue 302 to 390 of p53 (Klefstrom et al., 1997) and grown as a pool. gRNAs targeting mouse Il11 locus were designed using Benchling Software. (gRNA1: 5’ GCAGCCATTGTACATGCCGG3’ targeting exon 4, gRNA2: 5’AGGGGCAACGACTCTATCTG3’ targeting exon 2) and cloned into LentiCRISPRv2 plasmid as described (Sanjana et al., 2014; Shalem et al., 2014). For lentivirus production, 6 million HEK 293FT cells were seeded and transfected after 24 hours with the LentiCRISPRV2-IL11g1 or LentiCRISPRV2-IL11g2 construct together with the pLp Δ8,9 packaging plasmid and VSVG envelope plasmid using lipofectamine™ 2000 transfection reagent (Life Technologies, 11668019) and Opti-MEM reduced serum medium (Gibco, 31985-047). Viruses were collected after 48 hours, filtered through a 0.22 μm filter and stored at −80 °C until use. For transduction, 200.000 immortalized intestinal fibroblasts were seeded in 2 mL growth media on a 6-well plate. The media was replaced after 24 hours with 1 mL growth media mixed with 1 mL of virus media and 8 μg/mL of Polybrene (Merck Millipore, TR-1003-G). One day later, cells were split in growth media and selected using 2 μg/mL puromycin (Gibco, A1113803). The success of Il11 targeting was verified using Sanger sequencing.

### Quantitative PCR

qRT-PCR analysis was conducted to examine gene expression levels. For RNA extraction, the TRI Reagent (AM9738, Thermo Fisher Scientific) was used, following the manufacturer’s protocols. Subsequently, cDNA synthesis was performed using the TaqMan reverse transcription kit (N8080234, Applied Biosystems), and reverse transcription reactions were conducted using the MJ Research PTC-200 Thermal Cycler (Marshall Scientific). For qRT-PCR, the KAPA SYBR FAST qPCR Master Mix Universal reagent (KK4618, KAPA Biosystems) was utilized. All primers were diluted to a final concentration of 500 nM in nuclease-free water, with a reaction volume of 20 µl. StepOnePlus Real-Time PCR System (Thermo Fisher Scientific) was employed for sample analysis. Actb or Gapdh were used for normalization. The relative fold change in gene expression was determined using the ΔΔCT method described by Livak & (Livak & Schmittgen, 2001) (Livak & Schmittgen, 2001).

### scRNA-seq data analysis

Fastq files were aligned to the mouse reference genome (mm10), modified to include the TdTomato sequence, and demultiplexed using CellRanger.v3 (Zheng et al., 2017) setting a cell target number of 5000 cells per sample. Seurat V3 was used for further data processing. Pre-filtering was performed based on percent mitochondria (<15%), UMI count (<100000), and at least 400 features detected to exclude low-quality cells. SCT transformed setting *nfeatures* to 3000 was used for the selection of integration features, and further PrepSCTIntegration was used without reference sample. Based on an elbow plot, 30 principal components were used for RunUMAP and FindNeighbors functions. Clusters were identified using FindClusters, and for exploratory analysis, a range of different resolutions was used (0.3, 0.5, 0.7, 1.0, 1.3). Main clusters were manually mapped based on known markers (Figure 1, FigS1, S2). Data was further sub-clustered within the main cell category, taking the first cell cluster assignment and a further filter based on the expression of the primary marker genes: epithelial cells (Epcam >1 read), fibroblast (Pdpn >1 read), endothelial (Pecam >1 read), and immune (Ptprc >1 read). The subcluster dataset was re-analyzed using the same SCT transform-integration pipeline explained above. To assess the cell markers enriched in each cluster, the Seurat function FindAllMarkers was used with the default ‘roc’ method and a minimum percentage of cells threshold set at 0.2. For differential expression analysis of the overall differences between tumor and control, FindMarkers function was used with the method set to MAST (Finak et al., 2015) and *min.pct* of 0.2 with the active identities of the Seurat object set to be the tissue type (tumor vs control). ST2 signature of 25 genes was created by selecting the top significant up-regulated genes with a log fold-change above 1.5 against the rest of the fibroblast clusters (Supplementary Table 1).

### Cell-cell communication analysis

To analyze cell-cell communication, we utilized CellChat (Jin et al., 2021) with the following settings. A CellChat object was created from a Seurat object that included the identified clusters of all cells. This was achieved using the built-in function createCellChat from the CellChat R package. To ensure a comprehensive analysis, we utilized the complete CellChat ligand-receptor database, encompassing factors related to secretion, extracellular matrix (ECM), and cell-cell contact. We computed the communication probability and inferred the cellular communication network by executing the computeCommunProb method, wherein we set the parameters *raw.use* and *population.size* as False. This setting was chosen to prevent the size of the clusters from dominating the interaction strength in the dataset. Additionally, we applied a cell-cell communication filter with a minimum cell threshold of 10 (min.cells=10), thereby excluding communication pairs that accounted for only a few cells. For cell-cell communication pattern analysis, we used the CellChat built-in function *identifyCommunicationPatterns* with a rank of k=5 previously inferred by Cophenetic and Silhouette analysis as suggested in their documentation (https://github.com/sqjin/CellChat)

### Bulk RNA sequencing

RNA sequencing was performed using 3′UTR RNA sequencing with a ’Pool Seq’ method, which is based on the Dropseq approach for single-cell sequencing (Macosko et al., 2015). The method involves priming mRNA using an oligo-dT primer incorporating a 12 bp barcode and an 8 bp unique molecular identifier (UMI) sequence. The UMI enables the removal of PCR duplicates during data analysis. To generate double-stranded cDNA, the single-stranded cDNA was converted using the template-switch effect, followed by PCR amplification using SMART PCR primers. Subsequently, the samples were pooled, and PCR sequencing pools were created using Nextera i7 primers and the Dropseq P5 primers. Sequencing was performed on the NextSeq 500 (Illumina) using the NextSeq High Output 75 cycle flow cell. The obtained fastq files were aligned to the mouse reference genome (GRCm38) using the STAR-aligner (v2.7.8a) (Dobin et al., 2013). To demultiplex the samples and create a count matrix of UMI counts, the ’CreateDGEMatrix’ command from the BRB-seqToolsv1.4 suite of tools (https://github.com/DeplanckeLab/BRB-seqTools) was employed. The input for this analysis consisted of R2 aligned BAM files and R1 fastq files. The R1 reads had a length of 21 nucleotides and included a 12-nucleotide sample barcode, an 8-nucleotide UMI, and an additional nucleotide. For publication purposes, individual files for demultiplexed raw fastq files and the processed UMI count matrix of each sample, were generated using the demultiplex function from the BRB-seqTools suite. The CSC - IT Center for Science, Finland, provided the computational resources required for all the data processing steps. The Lkb1 deficient fibroblast signature was created by selecting significant genes upregulated above 2 log fold-change in Lkb1 KO (AdCre transduced) fibroblasts against wild-type fibroblasts (AdGFP transduced) (Supplementary Table 1).

## Data Availability

scRNA-seq dataset has been uploaded to ArrayExpress with accession number E-MTAB-13624, bulk RNA-seq files from primary intestinal fibroblasts from Lkb1fl/fl mice adenotransducted with AdCre or AdGFP and treated with or without LPS and have been upload it to ArrayExpress with the accession number E-MTAB-13622. Bulk RNA-seq files from colon tissue lysates of mice subjected to acute DSS colitis have been uploaded with the accession number E-MTAB-13631.

## Acknowledgments

This mouse work was carried out with the support of HiLIFE Laboratory Animal Centre Core Facility, University of Helsinki, Finland. The 3′ RNA sequencing analysis was conducted at the Biomedicum Functional Genomics Unit, located at the Helsinki Institute of Life Science and Biocenter, Finland, affiliated with the University of Helsinki. The gRNA constructs were generated by the Genome Biology Unit supported by HiLIFE and the Faculty of Medicine, University of Helsinki, and Biocenter Finland. LentiCRISPR v2 was a gift from Feng Zhang (Addgene plasmid #52961). Images were generated using 3DHISTECH Pannoramic 250 FLASH III digital slide scanner at the Genome Biology Unit supported by HiLIFE and the Faculty of Medicine, University of Helsinki, and Biocenter Finland. The flow cytometry was performed at the HiLIFE Flow Cytometry Unit, University of Helsinki. Roni Liimatta is thanked for help in gRNA design, and Melissa Montrose, Reeta Huhtala, Riikka Saikkonen and Myriam Sevigny for technical assistance. We are grateful to Chen Xie and Daryl Yeong for their technical assistance rendered to the colony maintenance, antibody treatments, and tissue harvests of the *Lkb1*^+/-^ mice performed at National Heart Centre Singapore, Singapore.

