## Supplementary Figures for "Identification of a targetable ST2-expressing fibroblast subset driving Peutz-Jeghers syndrome polyposis"

*Twist2-Cre;Lkb1*$^{flox/+}$, gastric polyp and adjacent normal antrum (control) scRNAseq

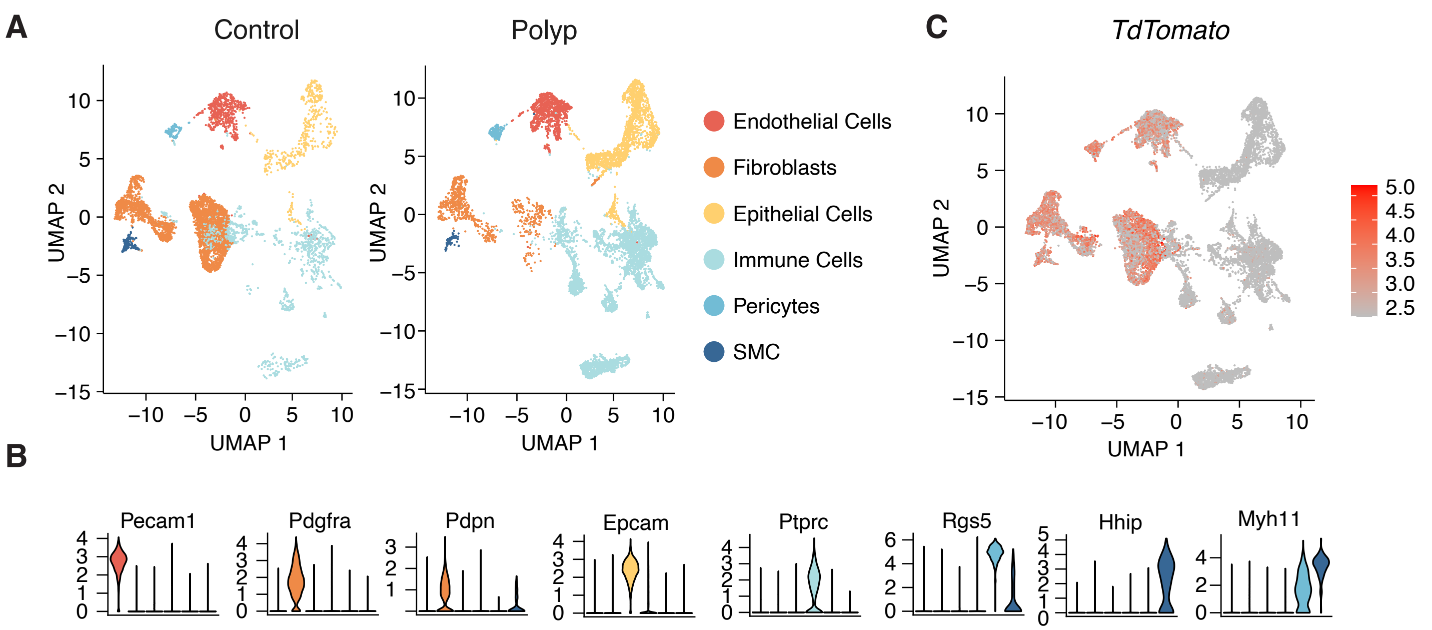


**Supplementary Figure 1. Single-cell sequencing of PJS type gastric tumors. (A)** UMAP clustering of all cells identified in the scRNAseq dataset from normal antrum (Control) and antral polyp. **(B)** Expression of tdTomato expression across the identified clusters. **(C)** Violin plots of the indicated marker genes as example markers used to identify the cell types.

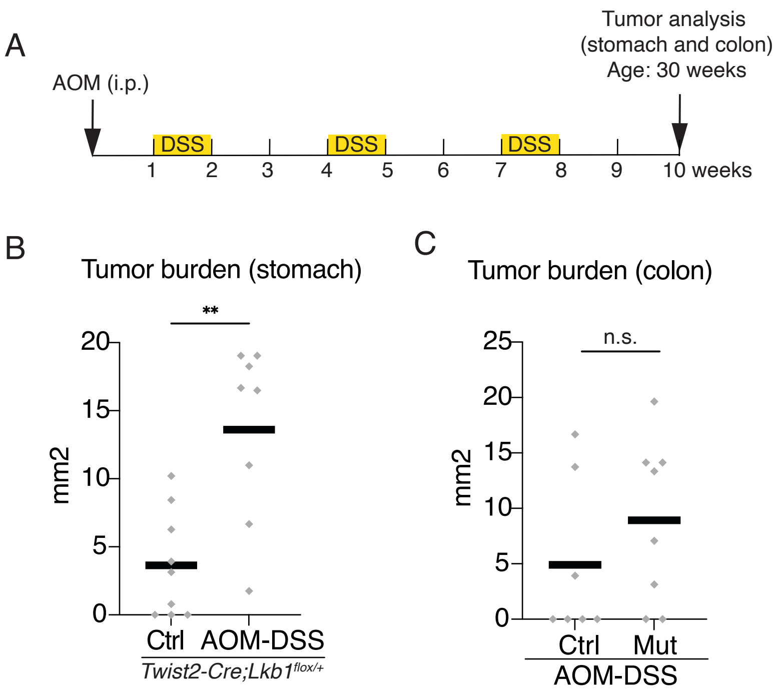


**Supplementary Figure 2. Inflammation leads to enhanced PJS polyposis. (A)** Outline of the inflammation-induced (AOM-DSS) model of colon tumorigenesis. AOM injection was performed at 20 weeks of age. **(B)** Gastric tumor burden in control or AOM-DSS treated Twist2-Cre;Lkb1fl/+ mice at 30 weeks of age. Ctrl, no treatment (n=9); AOM-DSS, n=8. **(C)** Colon tumor burden in control (Lkb1fl/+, n=7) and Mutant (Mut, Twist2-Cre;Lkb1fl/+, n=8) mice. **, p<0.01, Two-tailed T-test. n.s., not significant (p>0.05).

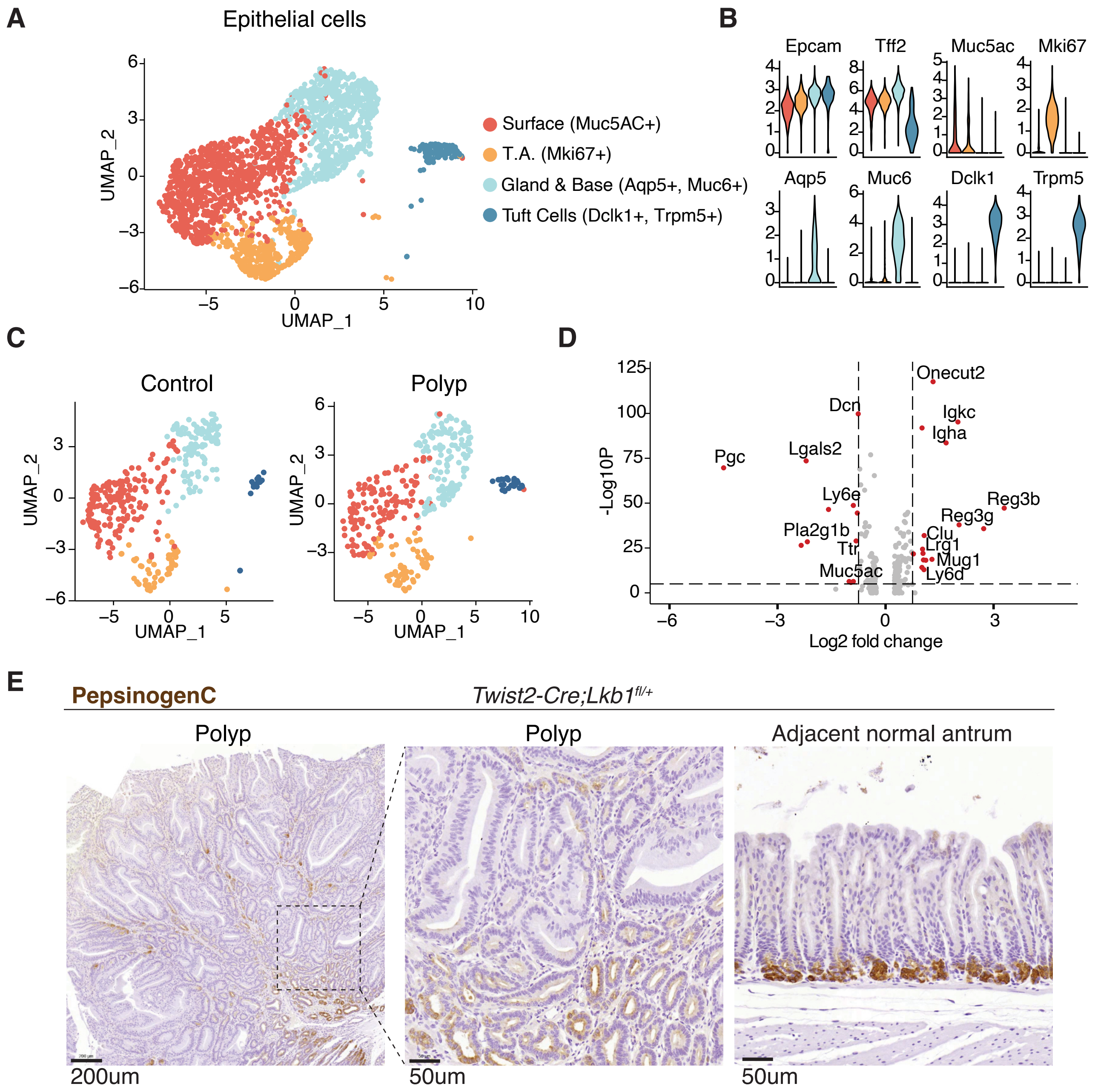


**Supplementary Figure 3. Decreased epithelial Pepsinogen C in the stroma-specific PJS model. (A)** UMAP clustering of all epithelial cells identified in the scRNAseq dataset. **(B)** Markers used to classify epithelial cell types. **(C)** UMAP clustering of the epithelial cell clusters in control antrum and polyp. Equal number of cells was plotted. **(D)** Volcano plot of differentially expressed genes between tumor cells and control antrum epithelial cells. Red dots depict significant genes with a log2 fold change above 1. **(E)** Immunohistochemical staining of Pepsinogen C in *Twist2-Cre;Lkb1fl/+* antral polyp sections and adjacent normal antrum.

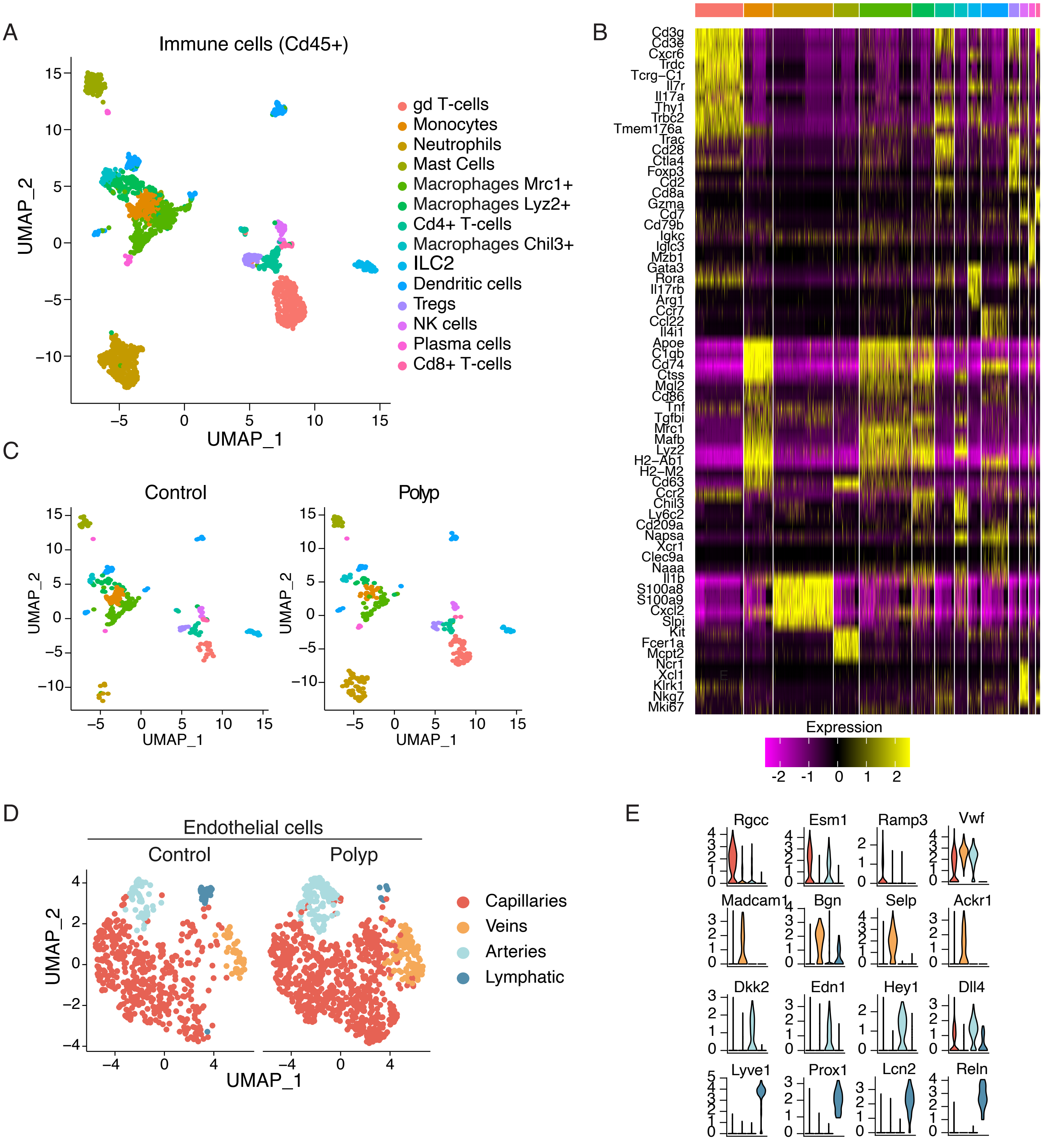


**Supplementary Figure 4. Analysis of immune cells and endothelial cells in gastric polyps. (A)** UMAP clustering of all immune cells identified in the scRNAseq dataset. **(B)** Markers used to classify the immune cell types. **(C)** UMAP clustering of the immune cell clusters in control antrum and polyp. Equal number of cells was plotted. **(D)** UMAP clustering of all endothelial cells identified in control antrum and polyp. **(E)** Markers used to classify the endothelial cell types.

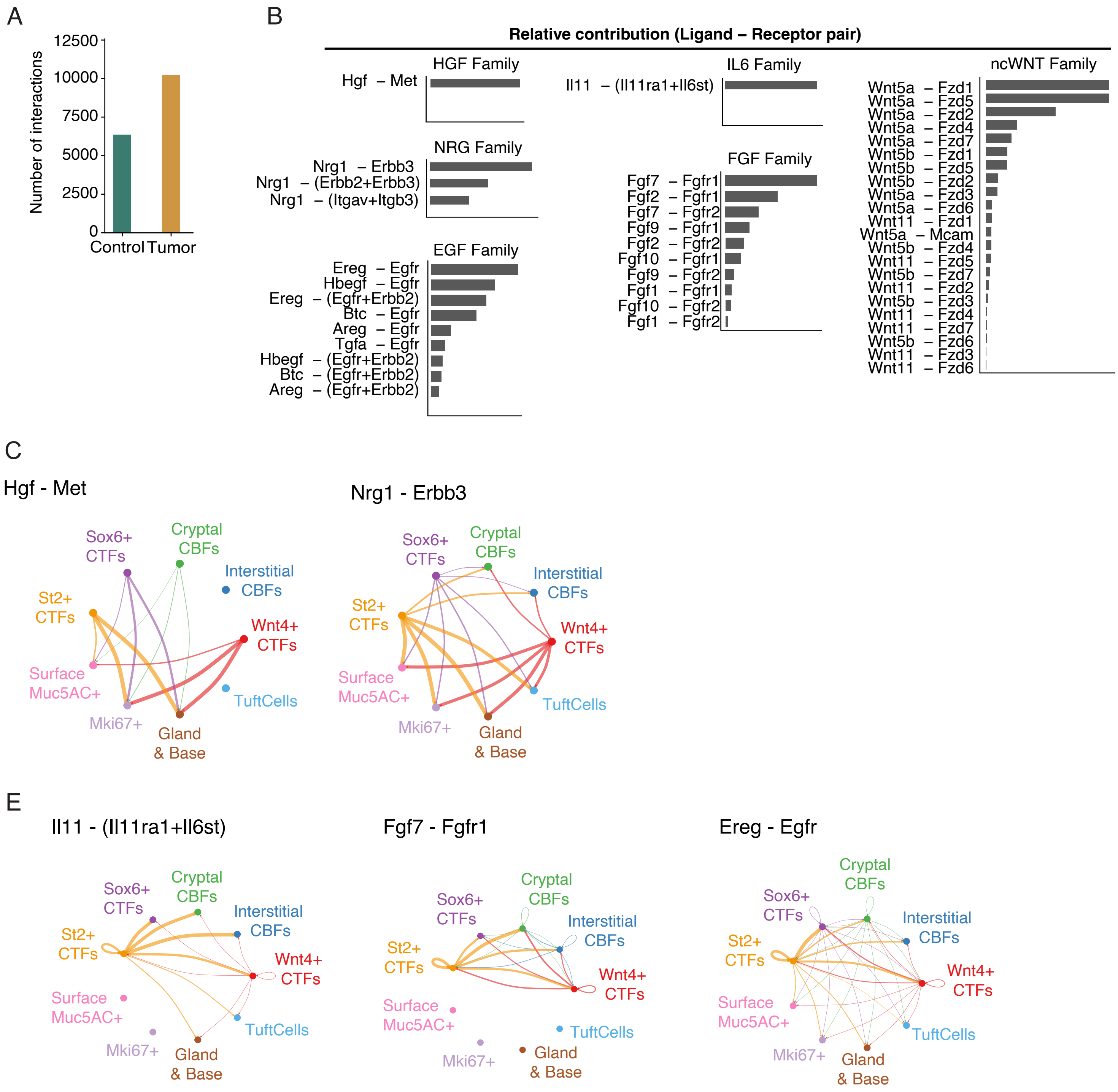


**Supplementary Figure 5. ST2-CTF-derived ligand-receptor signaling pairs. (A)** Total number of interactions within fibroblasts and epithelial cells in scRNAseq of control and polyp datasets. **(B)** Main ligand and receptor contributors for each family of ligands. **(C)** The most active ligand-receptor signaling interactions from ST2-CTFs to epithelial cell clusters. **(D)** The most active ligand-receptor signaling interactions from ST2-CTFs to other fibroblast clusters. **(C-D)** The line color represents the signaling source, and the line width the strength of the signaling. Only fibroblast and epithelial clusters were included in the circo plots.

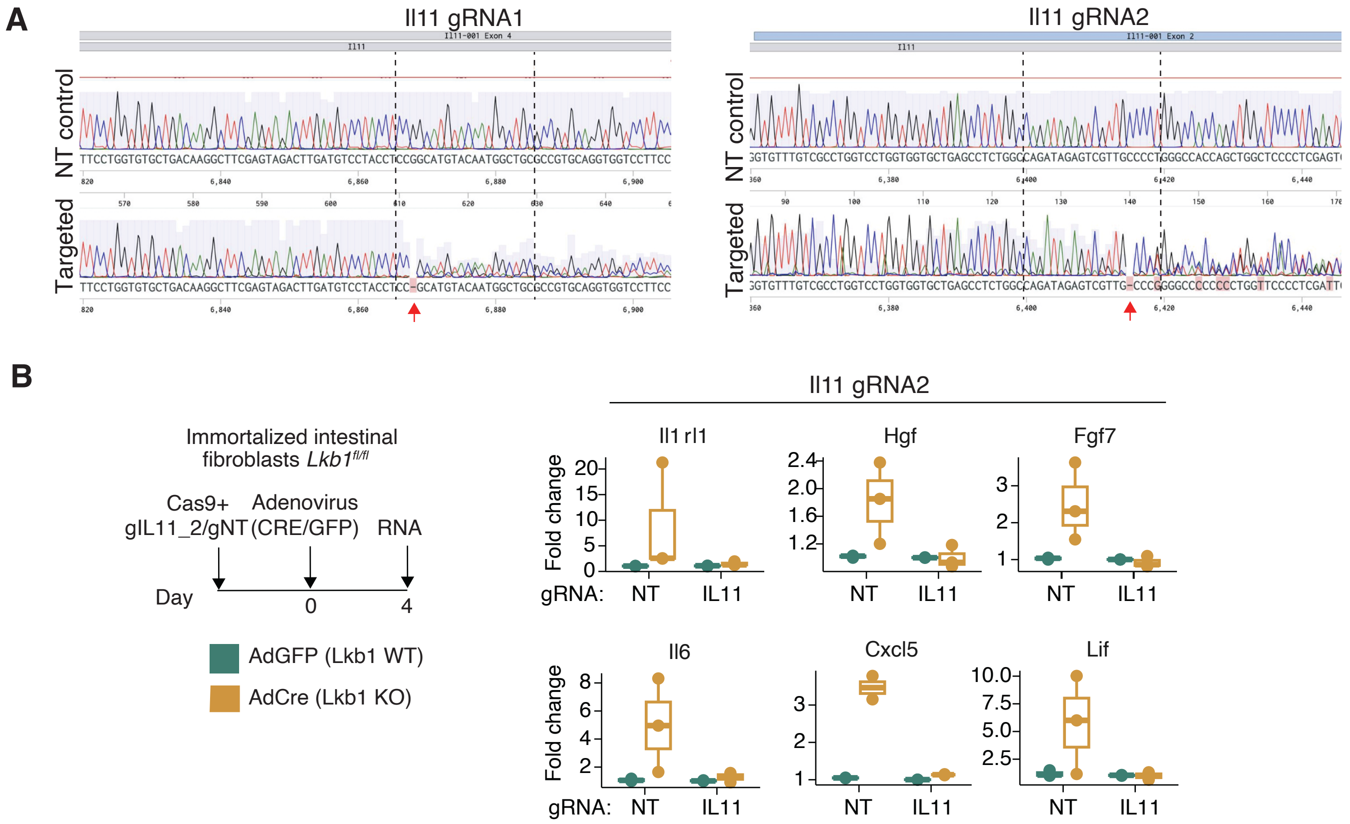


**Supplementary Figure 6. Generation of IL-11 deficient fibroblasts. (A)** Sanger sequencing of puromycin selected cell pools transduced with non-targeting (NT) gRNA or two independent gRNAs targeting Il11. The gRNA binding sites are highlighted, note mixed sequence and deletions (red arrows) in the targeted sequence. **(B)** Experimental setup (left) and gene expression of indicated genes in fibroblasts transduced with Cas9 and non-targeting (NT) gRNA or gRNA against IL11 (right). gRNA2 is shown, see also Figure 6C.
